# Phenome-wide Mendelian randomization mapping the influence of the plasma proteome on complex diseases

**DOI:** 10.1101/627398

**Authors:** Jie Zheng, Valeriia Haberland, Denis Baird, Venexia Walker, Philip Haycock, Mark Hurle, Alex Gutteridge, Pau Erola, Yi Liu, Shan Luo, Jamie Robinson, Tom G. Richardson, James R. Staley, Benjamin Elsworth, Stephen Burgess, Benjamin B. Sun, John Danesh, Heiko Runz, Joseph C. Maranville, Hannah M. Martin, James Yarmolinsky, Charles Laurin, Michael V. Holmes, Jimmy Liu, Karol Estrada, Rita Santos, Linda McCarthy, Dawn Waterworth, Matthew R. Nelson, Gibran Hemani, George Davey Smith, Adam S. Butterworth, Robert A. Scott, Tom R. Gaunt

## Abstract

The human proteome is a major source of therapeutic targets. Recent genetic association analyses of the plasma proteome enable systematic evaluation of the causal consequences of variation in plasma protein levels. Here, we estimated the effects of 1002 proteins on 225 phenotypes using two-sample Mendelian randomization (MR) and colocalization. Of 413 associations supported by evidence from MR, 130 (31.5%) were not supported by results of colocalization analyses, suggesting that genetic confounding due to linkage disequilibrium (LD) is widespread in naive phenome-wide association studies of proteins. Combining MR and colocalization evidence in cis-only analyses, we identified 111 putatively causal effects between 65 proteins and 52 disease-related phenotypes (www.epigraphdb.org/pqtl/). Evaluation of data from historic drug development programmes showed that target-indication pairs with MR and colocalization support were more likely to be approved, evidencing the value of our approach in identifying and prioritising potential therapeutic targets.

---

Despite increasing investment in research and development (R&D) in the pharmaceutical industry ^1^, the rate of success for novel drugs continues to fall ^2^. Lower success rates make new therapeutics more expensive, reducing availability of effective medicines and increasing healthcare costs. Indeed, only one in ten targets taken into clinical trials reaches approval ^2^, with many showing lack of efficacy (∼50%) or adverse safety profiles (∼25%) in late stage clinical trials after many years of development ^3 4^. For some diseases, such as Alzheimer’s disease, the failure rates are even higher ^5^.

Thus, early approaches to prioritize target-indication pairs that are more likely to be successful are much needed. It has previously been shown that target-indication pairs for which genetic associations link the target gene to related phenotypes are more likely to reach approval^6^. Consequently, systematically evaluating the genetic evidence in support of potential target-indication pairs is a potential strategy to prioritise development programmes. While systematic genetic studies have evaluated the putative causal role of both methylome and transcriptome on diseases ^7 8^, studies of the direct relevance of the proteome are in their infancy ^9 10^.

Plasma proteins play key roles in a range of biological processes, are frequently dysregulated in disease, and represent a major source of druggable targets ^11 12 13 14^. Recently published genome-wide association studies (GWAS) of plasma proteins have identified 3606 conditionally independent single nucleotide polymorphisms (SNPs) associated with 2656 proteins (‘protein quantitative trait loci’, pQTL) in more than 1000 participants ^9 15 16 17 18^. These genetic associations offer the opportunity to systematically test the causal effects of a large number of potential drug targets on the human disease phenome through Mendelian randomization (MR) ^19^. In essence, MR exploits the random allocation of genetic variants at conception and their associations with disease risk factors to uncover causal relationships between human phenotypes ^20^, and has been described in detail previously ^21 22^.

For MR analyses of molecular phenotypes such as proteins, unlike more complex exposures, an intuitive way to categorise protein-associated variants is into cis-acting pQTLs located in the vicinity of the encoding gene (defined as ≤ 500kb from the leading pQTL of the test protein in this study) and trans-acting pQTLs located outside this window. The cis-acting pQTLs are considered to have a higher biological prior and have been widely employed in relation to some phenome-wide scans of drug targets such as *CETP* ^23^, *HMGCR* ^12^, *PLA2G7* ^24^ and *IL6R* ^25 26^. Trans-acting pQTLs may operate via indirect mechanisms and are therefore more likely to be pleiotropic ^27 28^. However, non-pleiotropic trans-acting pQTLs may increase the reliability of the protein-phenotype associations.

Here, we pool and cross-validate pQTLs from five recently published GWAS and use them as instruments to systematically evaluate the potential causal role of 968 plasma proteins on the human phenome, including 153 diseases and 72 risk factors available in the MR-Base database ^29^. Results of all analyses are available in an open online database (www.epigraphdb.org/pqtl/), with a graphical interface to enable rapid and systematic queries.

## Results

### Characterising genetic instruments for proteins

**Figure 1** summarises the genetic instrument selection and validation process. We curated 3606 pQTLs associated with 2656 proteins from five GWAS ^9 15 16 17 18^. After removing proteins and SNPs in the major histocompatibility complex (MHC) region due to the complex LD structure of this region and performing strict LD clumping (r^2^<0.001 for SNPs in 10Mb window), we retained 2113 pQTLs associated with 1699 proteins and considered them as genetic instruments for the MR analysis (**Online Methods**: *Instrument selection*; instruments listed in **Supplementary Table 1**). Of these, 1062 of the 2113 instruments were identified in multiple studies (e.g. pQTLs from SOMAscan in Sun *et al*. ^9^ and pQTLs from OLINK in Folkersen *et al*.^16^) and the remaining 1051 pQTLs were only identified in one study. By testing the heterogeneity of the pQTLs effect across studies (**Online Methods:** *Consistency test estimating instrument heterogeneity across studies*), we found that the SNP effects of each of the pQTLs were similar across studies (78.5% of the pQTLs showed little evidence of heterogeneity) and effects of all pQTLs were also in high correlation across studies (**Supplementary Figure 1** and **2; Supplementary Table 2**). The proportion of cis and trans instruments in the two major platforms, SOMAscan and OLINK, were similar (SOMAscan: 38.1% cis and 61.9% trans instruments; OLINK: 44.8% cis and 55.2% trans instruments; **Supplementary Table 1**).

**Figure 1.**
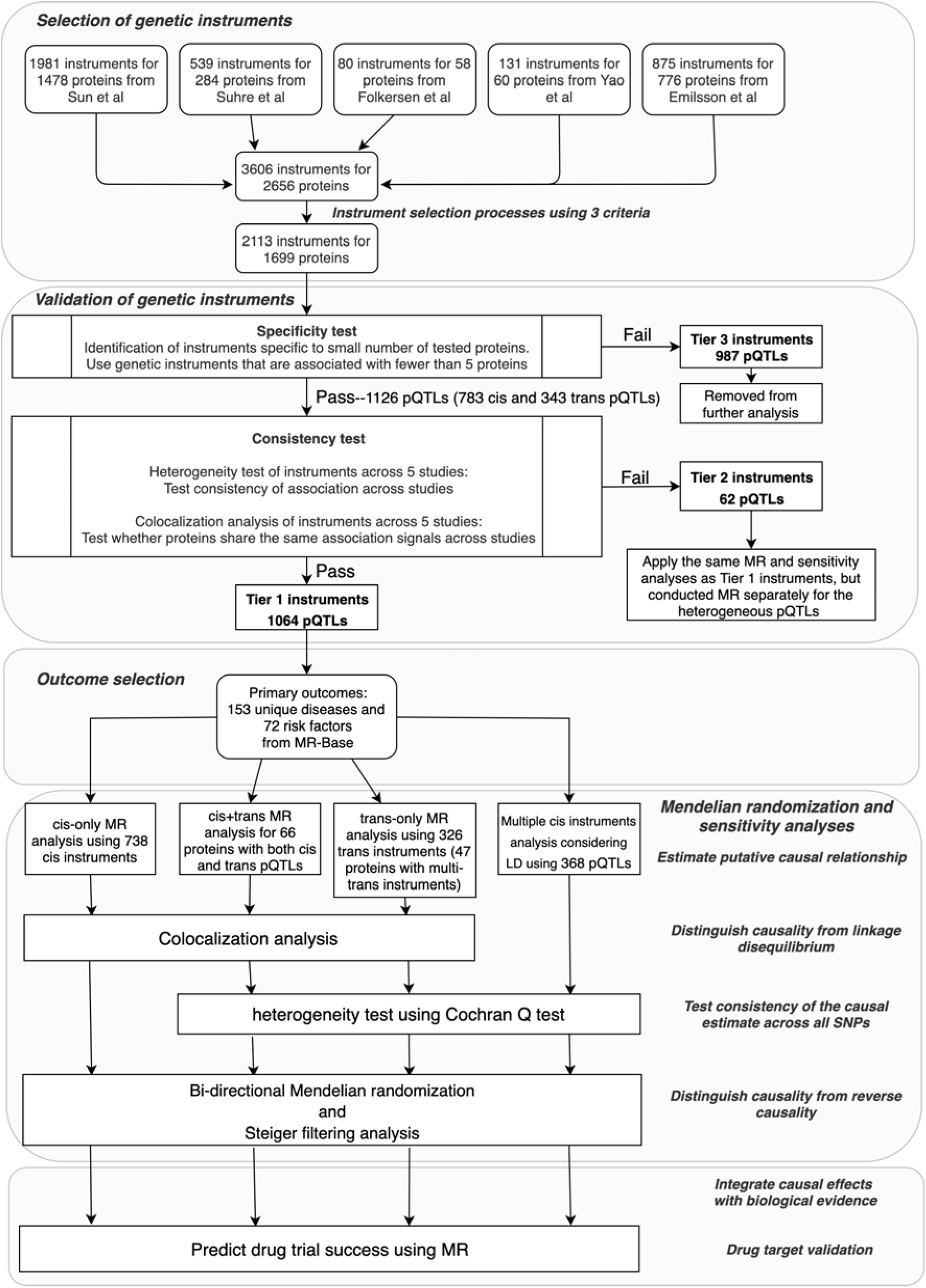
Study design of this phenome-wide MR study of the plasma proteome. The study included instrument selection and validation, outcome selection, 4 types of MR analyses, colocalization, sensitivity analyses and drug target validation.

We then conducted a validation process in which we categorised the instruments into three tiers based on their likely utility for MR analysis (**Online Methods**: *Instrument validation*). In summary, we curated 1064 instruments for 955 proteins with the highest relative level of reliability (tier 1, **Supplementary Table 1**), 62 instruments which exhibited SNP effect heterogeneity across studies (where we could test it), indicating uncertainty in the reliability of one or all instruments (tier 2, **Supplementary Table 1** and **3**), and 987 non-specific instruments which were associated with more than five proteins (tier 3, **Supplementary Table 1**). For the 263 tier 1 instruments associated with between two and five proteins, we aimed to distinguish between vertical and horizontal pleiotropy of these instruments by integrating protein-protein interaction (PPI) and pathway information (**Online methods**: *Distinguishing vertical and horizontal pleiotropic instruments using biological pathway data*). 68 instruments influenced multiple proteins in the same biological pathway (**Supplementary Table 1**), and thus are likely to reflect vertical pleiotropy and remain valid instruments since they do not violate the ‘exclusion restriction’ assumptions of MR ^22 27^.

Amongst the 1126 tier 1 and 2 instruments, 783 (69.5%) were cis-acting (within 500kb of the leading pQTL) and 343 were trans-acting. Of 1002 proteins with a valid instrument, 765 had only a single cis or trans instrument. 66 were influenced by both cis and trans SNPs (**Supplementary Table 4**) and 153 had multiple independent cis instruments (381 cis instruments showed in **Supplementary Table 5**).

### Estimated effects of plasma proteins on human phenotypes

We undertook two-sample MR to systematically evaluate evidence for the causal effects of 1002 plasma proteins (with tier 1 and tier 2 instruments) on 153 diseases and 72 disease related risk factors (**Supplementary Table 6, Online Methods**: *Phenotype selection*). As cis-pQTLs were considered to have a higher biological prior for a direct and specific impact of the SNP upon the protein (compared to trans pQTLs), we grouped the MR analyses based on whether the instruments were acting in cis or trans. Overall, we observed 413 protein-trait associations with MR evidence (P< 3.5×10^−7^ at a Bonferroni-corrected threshold) using either cis or trans instruments (or both for proteins with multiple instruments).

Genetically predicted associations between proteins and phenotypes may indicate causality (the protein causally influences the phenotype); reverse causality (genetic liability to a disease influences the protein); confounding by LD between the leading SNPs for proteins and phenotypes, or horizontal pleiotropy (the protein-phenotype association is not mediated by the target protein, but the dual associations are a result of two distinct biological phenomena) (**Supplementary Figure 3**). Given these alternative explanations, we conducted a set of sensitivity analyses designed to increase confidence that the MR association reflects a causal effect of the protein on the phenotype: tests of reverse causality using bi-directional MR ^30^ and MR Steiger filtering ^31 32^; heterogeneity analyses for proteins with multiple instruments ^33^, and colocalization analysis ^34^ to investigate whether the genetic associations with both protein and phenotype shared the same causal variant (**Figure 1**). To avoid unreliable inference from colocalization analysis due to the potential presence of multiple neighbouring association signals, we also developed and performed pair-wise conditional and colocalization analysis (PWCoCo) of all conditionally independent instruments against all conditionally independent association signals for the outcome phenotypes (**Online methods**: *Pair-wise conditional and colocalization analysis*; **Figure 2**).

**Figure 2.**
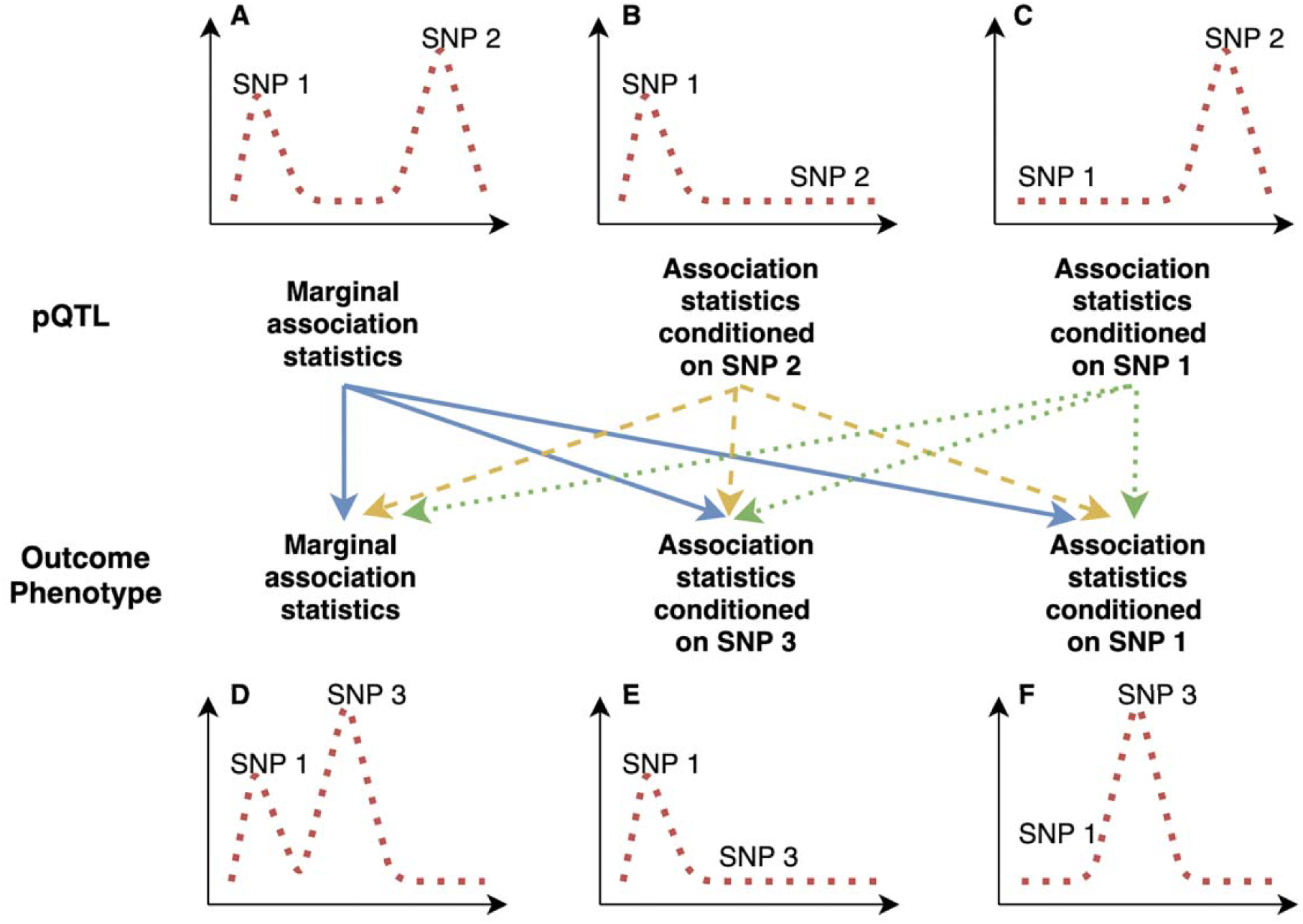
A demonstration of pair-wise conditional and colocalization (PWCoCo) analysis. Assume there are two conditional independent association pQTL signals (SNP 1 and SNP 2) and two conditional independent outcome signals (SNP 1 and SNP3) in the tested region. A naïve colocalization analysis using marginal association statistics will return weak evidence of colocalization (showed in regional plots A and D). By conducted the analyses conditioning on SNP 2 (plot B) and 1 (plot C) for the pQTLs and conditioning on SNP 1 (plot E) and 3 (plot F) for the outcome phenotype, each of the 9 pair-wise combinations of pQTL and outcome association statistics (represented as lines with different colours in the middle of this figure) will be tested using colocalization. In this case, the combination of plot B and plot E shows evidence of colocalization but the remaining 8 do not.

### Estimating protein effects on human phenotypes using cis pQTLs

In the cis-pQTL MR analyses, we identified 111 putatively causal effects of 65 proteins on 52 phenotypes (**Figure 3, Supplementary Table 7**), with strong evidence of MR (P< 3.5×10^−7^) and colocalization (posterior probability>80%; after applying PWCoCo) between the protein- and phenotype-associated signals. A further 69 potential associations had evidence from MR but did not have strong evidence of colocalization (posterior probability<80%; **Supplementary Table 8**), highlighting the potential for confounding by LD and the importance of colocalization analyses as a follow-up strategy in MR of proteins. Evidence of potentially causal effects supported by colocalization was identified across a range of disease categories including anthropometric and respiratory phenotypes, as well as cardiovascular and autoimmune diseases (**Supplementary Table 7**; **Supplementary Note 1**) and our findings replicated some previous reported associations (**Supplementary Note 2**).

**Figure 3.**
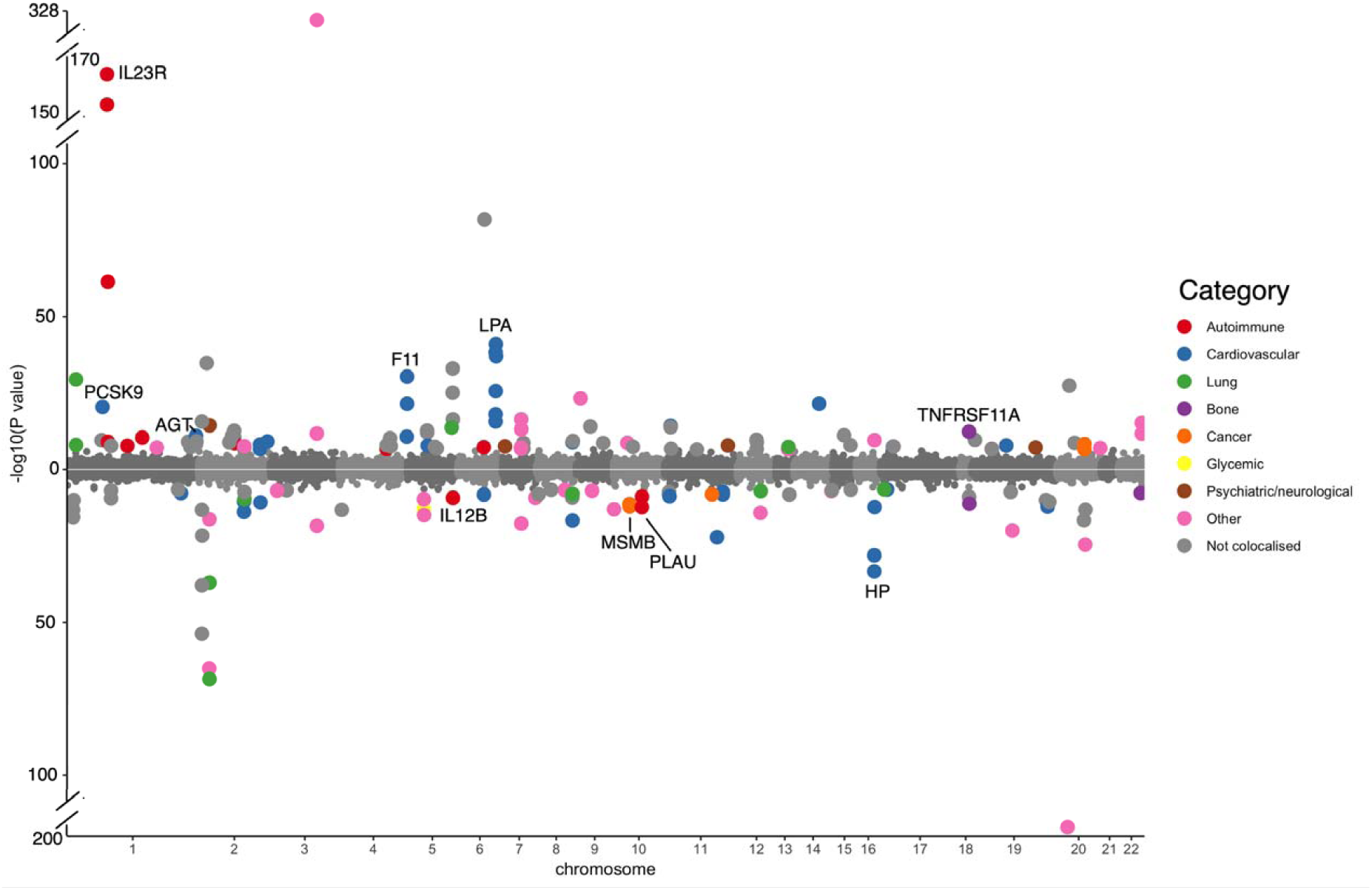
Miami plot for the cis-only analysis, with circles representing the MR results for proteins on human phenotypes. The labels refer to top MR findings with colocalization evidence, with each protein represented by one label. The colour refers to top MR findings with P<3.09×10^−7^, where red refers to immune mediated phenotypes, blue refers to cardiovascular phenotypes, green refers to lung related phenotypes, purple refers to bone phenotypes, orange refers to cancers, yellow refers to glycemic phenotypes, brown refers to psychiatric phenotypes, pink refers to other phenotypes and **grey** refers to phenotypes that showed less evidence of colocalization. The X-axis is the chromosome and position of each MR finding in the cis region. The Y-axis is the -log10 P value of the MR findings, MR findings with positive effects (increased level of proteins associated with increasing the phenotype level) are represented by filled circles on the top of the Miami plot, while MR findings with negative effects (decreased level of proteins associated with increasing the phenotype level) are on the bottom of the Miami plot.

Of 437 proteins with tier 1 or tier 2 cis instruments from Sun^9^ and Folkersen *et al*.^16^, 153 (35%) had multiple conditionally independent SNPs in the cis region identified by GCTA-COJO ^35^ (**Online methods**: *Instruments selection*; conditional signals in **Supplementary Table 5**). We applied an MR model which takes into account the LD structure between conditionally independent SNPs in these cis regions ^36 37^. In this analysis, we identified 10 additional associations, which had not reached our Bonferroni corrected P-value threshold in the single variant cis analysis (**Supplementary Table 9A**). Generally, the MR estimates from the multi-cis MR analyses were consistent with the single-cis instrumented analyses (**Supplementary Table 9B**).

In regions with multiple cis instruments, 16 of the 111 top cis MR associations only showed evidence of colocalization after conducting PWCoCo analysis for both the proteins and the human phenotypes (**Supplementary Table 7**). For example, interleukin 23 receptor (IL23R) had two conditionally independent cis instruments: rs11581607 and rs3762318 ^9^. Conventional MR analysis combining both instruments showed a strong association of IL23R with Crohn’s disease (OR=3.22, 95%CI= 2.93 to 3.53, P=6.93×10^−131^; **Supplementary Table 9B**). In addition, there were 4 conditionally independent signals (conditional P value<1×10^−7^) predicted for Crohn’s disease in the same region (Crohn’s disease data from de Lange *et al* 38). In the marginal colocalization analyses, we observed no evidence of colocalization (**Figure 4** and **Supplementary Figure 4**, colocalization probability=0). After performing PWCoCo with each distinct signal in an iterative fashion, we observed compelling evidence of colocalization between IL23R and Crohn’s disease for the top IL23R signal (rs11581607) (**Figure 4**, colocalization probability=99.3%), but limited evidence for the second conditionally independent IL23R hit (rs7528804) (colocalization probability = 62.9%). Additionally, for haptoglobin, which showed MR evidence for LDL-cholesterol (LDL-C), there were two independent cis instruments. There was little evidence of colocalization between the two using marginal associations (colocalization probability=0.0%). However, upon performing PWCoCo, we observed strong evidence of colocalization for both instruments (colocalization probabilities = 99%; **Supplementary Table 10; Supplementary Figure 5**). Both examples demonstrate the complexity of the associations in regions with multiple independent signals and the importance of applying appropriate colocalization methods in these regions. Of the 413 associations with MR evidence (using cis and trans instruments), 283 (68.5%) also showed strong evidence of colocalization using either a traditional colocalization approach (260 associations) or after applying PWCoCo (23 associations), suggesting that one third of the MR findings could be driven by genetic confounding by LD between pQTLs and other causal SNPs.

**Figure 4.**
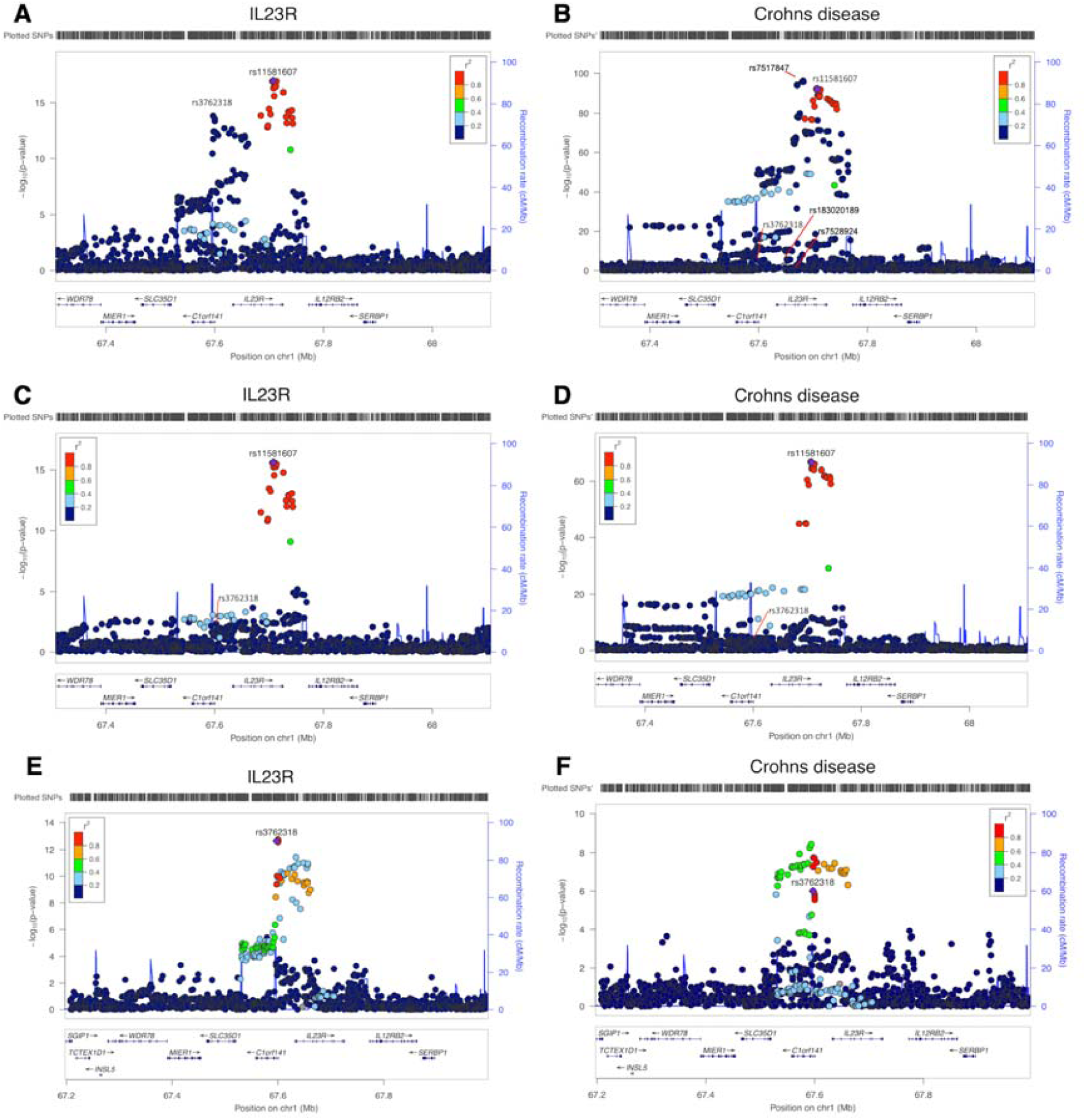
Regional association plots of IL23R plasma protein level and Crohn’s disease in the IL23R region. A. and B. the regional plots of IL23R protein level and Crohn’s disease without conditional analysis, Plot B listed the sets of conditional independent signals for Crohn’s disease in this region: rs7517847, rs7528924, rs183020189, rs7528804 (a proxy for the second IL23R hit rs3762318, r2=0.42 in the 1000 Genome Europeans) and rs11209026 (a proxy for the top IL23R hit rs11581607, r2=1 in the 1000 Genome European), conditional P value < 1×10-7; C. the regional plot of IL23R with the joint SNP effects conditioned on the second hit (rs3762318) for IL23R; D. the regional plot of Crohn’s disease with the joint SNP effects adjusted for other independent signals except top IL23R signal rs11581607; E. the regional plot of IL23R with the joint SNP effects conditioned on the top hit (rs11581607) for IL23R; F. the regional plot of Crohn’s disease with the joint SNP effects adjusted for other independent signals except second IL23R signal rs3762318. The heatmap of the colocalization evidence for IL23R association on Crohn’s disease (CD) in the IL23R region was presented in **Supplementary Figure 4**.

Due to potential epitope-binding artefacts driven by protein-altering variants, some of the cis instruments could be artefactual ^39^. We therefore conducted a sensitivity MR analysis that excluded 122 tier 1 or tier 2 cis instruments that are in the coding region or in LD (r^2^>0.8) with genetic variants in the coding region (**Supplementary Table 7 and 8**, filtered by column “VEP” including missense, stop-lost/gained, start-lost/gained and splice-altering variants). Of the 538 cis instruments for which we were still able to conduct an MR analysis after removing variants in the coding region, 77 protein-phenotype associations still had strong MR and colocalization evidence, suggesting that these are robust to epitope-binding artefacts.

### Using trans-pQTLs as additional instrument sources

Trans pQTLs are more likely to influence targets though pleiotropic pathways. For example, among the 1316 trans instruments we identified from 5 studies, 73.5% were associated with more than 5 proteins, compared with 1.8 % of cis instruments (**Supplementary Table 1**). However, trans pQTLs that overlap disease associations can highlight previously unsuspected candidate proteins through which loci may influence disease risk ^9^. In a MR context, including non-pleiotropic trans-pQTLs may increase the reliability of the protein-phenotype associations since (1) they will increase variance explained of the tested protein and increase power of the MR analysis; (2) the causal estimate will not be reliant on a single locus; and (3) further sensitivity analyses, such as heterogeneity test of MR estimates across multiple instruments, can be conducted. Therefore, we extended our MR analyses to include 343 trans instruments, where they were associated with fewer than 5 proteins. 192 of the 343 (56%) trans instruments were associated with just one protein or involved in a single biological pathway (**Supplementary Figure 6B**).

We implemented two strategies to utilize trans instruments to increase power of identifying causal links between proteins and phenotypes. Firstly, we combined cis and trans instruments for 66 proteins that had both cis and trans instruments (noted as cis + trans analysis). However, none reached our pre-defined Bonferroni-corrected threshold, and only two protein-phenotype associations showed some weak evidence (P<1×10^−5^) (**Supplementary Table 11**). Secondly, we performed trans-only MR analyses of 293 proteins, and identified 158 associations with 44 phenotypes that also had strong evidence (posterior probability>0.8) of colocalization (**Supplementary Table 12**). A further 54 trans-only MR associations did not have strong evidence of colocalization (**Supplementary Table 13**).

Some of the trans analyses with MR and colocalization evidence suggest causal pathways that are confirmed by evidence from rare pathogenic variants or existing therapies. For example, although we had no cis instrument for Protein C (Inactivator Of Coagulation Factors Va And VIIIa) (PROC) (**Supplementary Figure 7A**), we found strong evidence for a causal association between PROC levels and deep venous thrombosis (DVT) (P=1.27×10^−10^; colocalization probability>0.9) using a trans pQTL (rs867186; **Supplementary Figure 7B**), which is a missense variant in *PROCR* ^40^, the gene encoding the endothelial protein C receptor (EPCR). Patients with mutations in *PROC* have protein C deficiency, a condition characterised by recurrent venous thrombosis for which replacement protein C is an effective therapy.

From 47 proteins with multiple trans instruments, we identified four additional MR associations, but none showed strong evidence of colocalization (**Supplementary Table 13**). Proteins with multiple trans instruments showed little evidence of heterogeneity, which is comparable to the multiple cis MR (**Supplementary Table 14**).

### Estimating protein effects on human phenotypes using pQTLs with heterogeneous effects across studies

Among the 2113 selected instruments in **Supplementary Table 1**, we checked whether the same instrument was observed in other studies. We found 50.2% (1062) of the instruments (or proxy SNPs with r^2^>0.8) had association information in at least two studies (**Supplementary Table 15**). For these 1062 SNPs, we examined any differences in effect size between studies using the pair-wise Z test (where we defined a Z statistic greater than 5 as indicating evidence for heterogeneity). Of the 494 tier 1 or tier 2 instruments where we could test for heterogeneity across studies, we found that 62 (12.6%) showed evidence of difference in effect size across studies (so called Tier 2 instruments). Recognising that effect heterogeneity does not preclude identification of genuine causal effects, we performed MR analyses using the most significant SNP across studies and report the findings with caution. Some proteins that are targets of approved drugs were found to have potential causal effects in this analysis, such as interleukin-6 receptor (IL6R) on rheumatoid arthritis ^41^, and coronary heart disease (CHD) ^25 26^ (**Supplementary Table 16**). Tocilizumab, a monoclonal antibody against IL6R, is used to treat rheumatoid arthritis, while canakinumab, a monoclonal antibody against interleukin-1 beta (an upstream inducer of interleukin-6), has been shown to reduce cardiovascular events specifically among patients who showed reductions in interleukin-6 ^42^.

As another test of heterogeneity across studies, where the same protein was measured in two or more studies, we performed colocalization analysis of each pQTL (in one study) against the same pQTL (in another study) for the two studies in which we had access to full summary results (Sun *et al*. ^9^ and Folkersen *et al*. ^. 16^). Of the 41 proteins measured in both studies, 78 pQTLs could be tested using conventional colocalization and PWCoCo (**Supplementary Table 15**). We found weak evidence of colocalization for 51 pQTLs (posterior probability<0.8), which suggested either two different signals were present within the test region or the protein has a pQTL in one study but not in the other. In either case, as one of the two distinct signals may be genuine, we performed MR analysis of these 25 pQTLs using instruments from each study separately. 8 associations had MR evidence but only one showed colocalization evidence (IL27 levels on human height; **Supplementary Table 17**).

### Orienting causal direction in protein-phenotype associations

For potential associations between proteins and phenotypes identified in the previous analyses (single cis, cis + trans and trans-only analyses), we undertook two sensitivity analyses to highlight results due to reverse causation: bi-directional MR ^30^ and Steiger filtering ^31 32^ (**Online Methods**: *Distinguishing causal effects from reverse causality*). In general, we found no strong evidence of reverse causality for genetic predisposition to diseases on protein level changes, in either bi-directional MR analyses or Steiger filtering. Of 360 associations to which we were able to apply the MR Steiger filtering analysis, only 57 (15.8%) showed some evidence of reverse causality (i.e. effect from human phenotypes to proteins), with the majority of these being trans instruments (n=56) (**Supplementary Note 3** and **Supplementary Data 1**).

### Drug target prioritisation and repositioning using phenome-wide MR

Recent MR studies highlight the value of hypothesis-free (“phenome-wide”) MR in building a comprehensive picture of the causal effects of risk factors on human phenotypes ^8 43 44^. Given that human proteins represent the major source of therapeutic targets, we sought to mine our results for targets of molecules already approved as treatments or in ongoing clinical development. We first compared MR findings for 1002 proteins against 225 phenotypes with historic data on progression of target-indication pairs in Citeline’s PharmaProjects (downloaded on the 9^th^ of May 2018). Of 783 target-indication pairs with an instrument for the protein and association information for a phenotype similar to the indication for which the drug had been trialled, 9.2% (73 pairs) had successful (approved) drugs, 69.1% had failed drugs (including 195 drugs which in the clinical stage and 354 drugs failed in the preclinical stage) and 20.3% were for drugs still under development (161 pairs). The 268 pairs for successful (73) or failed (195) drugs (88 unique proteins and 66 unique phenotypes) were selected for further analyses (**Supplementary Table 19**). We observed 8 target-indication pairs for successful drugs with positive MR and colocalization evidence (**Supplementary Table 20**). In addition to the PROC and IL6R examples discussed earlier, we found Proprotein convertase subtilisin/kexin type 9 (PCSK9) (target for evolocumab) for hypercholesterolemia and hyperlipidemia, Angiotensinogen (AGT) for hypertension, IL12B for psoriatic arthritis and psoriasis and TNF Receptor Superfamily Member 11a (TNFRSF11A) for osteoporosis. Of 195 target indication pairs that had failed to gain approval after clinical trials for a variety of reasons, none had sufficient MR and colocalization evidence to reach our threshold. We further removed 75 target indication pairs which may upwardly bias the comparison between MR and drug trials from the 268 pairs (**Online Methods**: *Drug target validation and repositioning*) (**Supplementary Table 21**). The comparison using the remaining 193 pairs indicated that protein-phenotype associations with MR and colocalization evidence are more likely to be successful drugs (**Table 1**). Although we acknowledge the limited sample size of the test set, this does support the utility of pQTL MR analyses with colocalization as a source of target identification and prioritization.

**Table 1.**
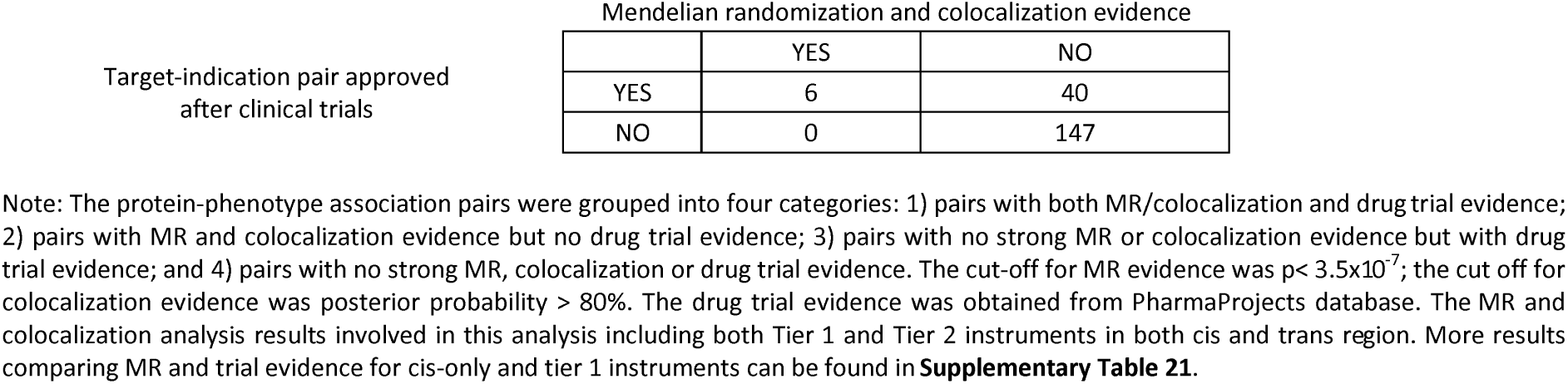
Enrichment analysis comparing target-indication pairs with or without MR and colocalization evidence.

Previous efforts have highlighted the opportunity of using genetics for drug repositioning ^45^. We identified 3 approved drugs for which we found pQTL MR and colocalization evidence for 5 phenotypes other than the primary indication and 23 drug targets under development for 33 alternative phenotypes (**Supplementary Table 22**). For example, our phenome-wide MR analysis suggested that lifelong higher urokinase-type plasminogen activator (PLAU) levels are associated with lower inflammatory bowel disease (IBD) risk (OR=0.75, 95%CI= 0.69 to 0.83, P= 1.28×10^−9^; **Supplementary Figure 8**), potentially identifying a repositioning opportunity for IBD. However, we note this opportunity with caution given the multitude of considerations in such a strategy. For example, the drug Kinlytic (urokinase) was initially developed for use as a thrombolytic in the treatment of acute myocardial infarction and ischaemic stroke, and thus a target-mediated adverse effect is an increase in bleeding and potential haemorrhage. In addition, the current agent is administered intravenously which precludes it as an option as a long-term preventative treatment. While our data suggest that Kinlytic might be protective in the aetiology of IBD, a careful risk benefit assessment would be required as part of an investigation into whether drugs targeting urokinase might be repurposed for the treatment of IBD.

We also evaluated drugs in current clinical trials and identified 8 additional protein-phenotype associations with MR and colocalization evidence. Examples include lipoprotein(a) (LPA) for blood lipids and angiopoietin like 3 (ANGPTL3) for blood lipids (**Supplementary Table 23**), for which we observe MR evidence implicating an increased likelihood of success.

Finally, we compared the 1002 instrumentable proteins (i.e. those that passed our instrument selection procedure) against the druggable genome ^46^. 682 of the 1002 instrumentable proteins overlapped with the druggable genome (all proteins: 68.1%; proteins with cis instruments: 72.6%; proteins with trans instruments: 60.1%; **Supplementary Table 24** and **Online Methods**: *Enrichment of proteome-wide MR with the druggable genome*). A further enrichment analysis was conducted to assess the overlap between putative causal protein-phenotype associations and the druggable genome (**Supplementary Table 25**). Of the 295 top findings (120 proteins on 70 phenotypes) with both MR and colocalization evidence, 250 of them (87.7%) overlapped with the druggable genome (**Figure 5** and **Supplementary Table 25**). Although the druggable genome is continuously evolving ^47 48 49 50^, this enrichment analysis reveals that MR findings for the tested bone diseases, cancers and psychiatric diseases are less represented in tier 1 of the druggable genome, in comparison with cardiovascular and immune-mediated diseases.

**Figure 5.**
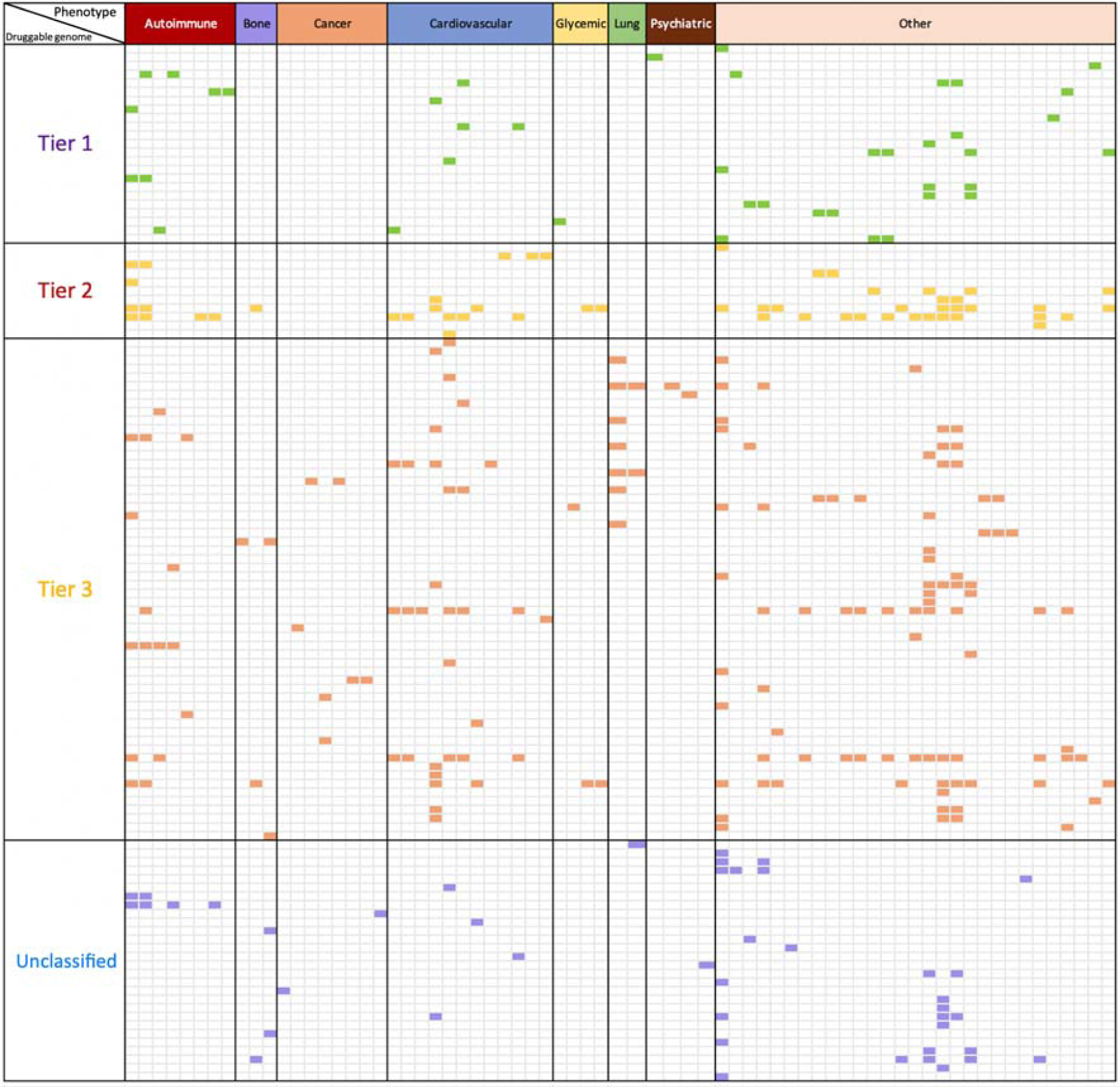
Enrichment of phenome-wide MR of the plasma proteome with the druggable genome. In this figure, we only showed proteins with convincing MR and colocalization evidence with at least one of the 70 phenotypes. The X-axis shows the categories of 70 human phenotypes, where the phenotypes have been grouped into 8 categories: 8 autoimmune diseases (red), 3 bone phenotypes (purple), 8 cancers (orange), 12 cardiovascular phenotypes (blue), 4 glycemic phenotypes (yellow), 2 lung phenotypes (green), 4 psychiatric phenotypes (brown) and 29 other phenotypes (pink). The Y-axis presents the tiers of the druggable genome (as defined by Finan *et al*) of 120 proteins under analysis, where the proteins have been classified into 4 groups based on their druggability: tier 1 contained 23 proteins which are efficacy targets of approved small molecules and biotherapeutic drugs, tier 2 contained 11 proteins closely related to approved drug targets or with associated drug-like compounds, tier 3 contained 58 secreted or extracellular proteins or proteins distantly related to approved drug targets, and 28 proteins have unknown druggable status (Unclassified). The cells with colours are protein-phenotype associations with strong MR and colocalization evidence. Cells in green are associations overlapped with tier 1 druggable genome, where cells in yellow, red or purple were associations with tier 2, tier 3 or unclassified. More detailed information shown in **Supplementary Table 25**.

## Discussion

MR analysis of molecular phenotypes against disease phenotypes provides a promising opportunity to validate and prioritise novel or existing drug targets through prediction of efficacy and potential on-target beneficial or adverse effects ^51 52 53^. Our phenome-wide MR study of the plasma proteome employed five pQTL studies to robustly identify and validate genetic instruments for thousands of proteins. We used these instruments to evaluate the potential effects of modifying protein levels on hundreds of complex phenotypes available in MR-Base ^29^ in a hypothesis-free approach ^19^. We confirmed that protein-phenotype associations with both MR and colocalization evidence predicted a higher likelihood of a particular target-indication pair being successful and highlight 283 potentially causal associations (111 in the cis-only analysis). Collectively, we underline the important role of pQTL MR analyses as an evidence source to support drug discovery and development and highlight a number of key analytical approaches to support such inference.

In particular, we note the distinct opportunities and requirements for MR of molecular phenotypes, such as proteins, compared to other complex exposures such as BMI ^54^. Particular features of such molecular exposures require a different approach. For example, the number of instruments is often limited, restricting the opportunity to apply recently developed pleiotropy robust approaches ^55 56 33^. New methods such as MR-robust adjusted profile scoring (MR-RAPS) ^57^ allow inclusion of many weak instruments in the MR analysis and have been applied to a recent proteome-wide MR study ^10^. However, we note some examples where inclusion of multiple weaker instruments can reduce power and yield different results to those based on cis instruments alone ^51 58^. A major advantage of proximal molecular exposures such as transcriptomics and proteomics is the ability to include cis instruments (or interpretable trans instruments) with high biological plausibility, limiting the likelihood of horizontal pleiotropy ^27 28^. However, undue focus on single SNP MR approaches brings susceptibility to other pitfalls ^22^. For example, our understanding of the effects of IL6R and PROC on the soluble protein levels is not enough to make strong inference of the direction of effects.

To provide robust MR estimates for proteins, we note the important role of a number of sensitivity analyses following the initial MR in order to distinguish causal effects of proteins from those driven by horizontal pleiotropy, genetic confounding through LD ^22^ and/or reverse causation ^31 32^. Of note, only two-thirds of our putative causal associations had strong evidence of colocalization, suggesting that a substantial proportion of the initial findings were likely to be driven by genetic confounding through LD between pQTLs and other disease-causal SNPs. To avoid misleading results, we suggest that for regions with multiple molecular trait QTLs, it is important to consider methods such as PWCoCo, which can avoid the assumptions of traditional colocalization approaches of just a single association signal per region ^59^. In the current study, application of PWCoCo identified evidence of colocalization for 23 additional protein-phenotype associations hidden to marginal colocalization ^59^. We note the proliferation of approaches to identify shared genetic effects between molecular phenotypes and disease ^60 63 64^, and that recent recommendations support the use of colocalization as a follow up analysis to reduce false positives ^63 64^.

An important limitation of this work is that protein levels are known to differ between cell types ^65^. In this study, we have estimated the role of protein measured in plasma on a range of complex human phenotypes but are unable to assess the relevance of protein levels in other tissues. Whilst eQTL studies highlight a large proportion of eQTLs being shared across tissues ^37^, there are many which show cell type and state specificity ^66^, highlighting the potential value of applying the current approach to data from proteomics analyses in other cell types and tissues. We also hypothesize that in instances with multiple conditionally distinct pQTLs, but where we observe colocalization of only certain conditionally distinct pQTL-phenotype pairs, that this may reflect underlying cell- and state-specific heterogeneity in bulk plasma pQTLs, among which only certain cell-types or states are causal ^67^. Although pQTL studies have not yet been performed as systematically across tissues or states as eQTL studies, it remains encouraging that our analyses using plasma proteins identify associations across a range of disease categories, including for psychiatric diseases for which we may expect key proteins to function primarily in the brain.

Evaluating the potential of MR to inform drug target prioritisation, we demonstrated that the presence of pQTL MR and colocalization evidence for a target-indication pair predicts a higher likelihood of approval. One of the limitations of our approach is the lack of comprehensive coverage of genetic data for all phenotypes for which drugs are in development, as well as our inability to instrument the entire proteome through pQTLs. As such, ongoing expansions in the scale, diversity and availability of GWAS will be important in providing more precise estimates of the value of MR and colocalization in drug target prioritization and in enabling its broader application.

Another potential limitation of our work is the presence of epitope-binding artefacts driven by coding variants that may yield artefactual cis pQTLs ^39^. In particular, such instances may lead to false negative conclusions where, in the presence of a silent missense variant causing an artefactual pQTL but with no actual effect on protein function or levels, we do not correctly instrument the target protein. In instances where the missense variant appears to be driving the association with the phenotype, we suggest that causal inference may remain valid but inference on direction of association is challenged. Finally, the limited coverage of the proteome afforded by current technologies, leaves the possibility of undetected pleiotropy of instruments. While cis-pQTLs are less likely to be prone to horizontal pleiotropy than trans-pQTLs, it is well known from studies of gene expression that cis variants can influence levels of multiple neighbouring genes and hence the same is likely to be true for proteins ^68^. Future larger GWAS of the plasma proteome are likely to uncover many more variant-protein associations, increasing the apparent pleiotropy of many pQTLs.

## Conclusion

In conclusion, this study systematically identified 283 putatively causal effects between the plasma proteome and the human phenome using the principles of MR and colocalization. These observations support, but do not prove, causality, as potential horizontal pleiotropy remains an alternative explanation. Our study provides both an analytical framework and an open resource to prioritise potential new targets on the basis of MR evidence and a valuable resource for evaluation of both efficacy and repurposing opportunities by phenome-wide evaluation of on-target associations.

## Author contribution

JZ, VH and DB performed the Mendelian randomization analysis; JZ and DB performed the colocalization analysis; JZ performed the conditional analysis; VH, YL, BE and TRG developed the database and web browser; JZ and VW performed the drug target prioritisation and enrichment analysis. JZ and RS conducted the druggable genome analysis; JZ and PE conducted the pathway and protein-protein interaction analysis; AG, TGR, BE, HM, JY, CL, SL and JR conducted supporting analyses; JS, BBS, JD, HR, JCM provided key data and supported the MR analysis; JL, KE, LM, MVH, MH, DW, MRN reviewed the paper and provided key comments. JZ, VH, DB, VW, PH, AB, GDS, GH, RAS and TRG wrote the manuscript. JZ, TRG and RAS conceived and designed the study and oversaw all analyses.

## Acknowledgements

We are extremely grateful to all the families who took part in the ALSPAC study, the midwives for their help in recruiting them, and the whole ALSPAC team, which includes interviewers, computer and laboratory technicians, clerical workers, research scientists, volunteers, managers, receptionists and nurses. We acknowledge Jack Bowden for statistical support and advice relating to MR-Egger regression.

This publication is the work of the authors and Jie Zheng will serve as guarantor for the contents of this paper. JZ is funded by a Vice-Chancellor Fellowship from the University of Bristol. This research was also funded by the UK Medical Research Council Integrative Epidemiology Unit (MC_UU_00011/1 and MC_UU_00011/4), GlaxoSmithKline, Biogen and the Cancer Research Integrative Cancer Epidemiology Programme (C18281/A19169). The UK Medical Research Council and Wellcome (Grant ref: 102215/2/13/2) and the University of Bristol provide core support for ALSPAC. A comprehensive list of grants funding is available on the ALSPAC website (http://www.bristol.ac.uk/alspac/external/documents/grant-acknowledgements.pdf). GH is funded by the Wellcome Trust and the Royal Society [208806/Z/17/Z]. MVH is supported by a British Heart Foundation Intermediate Clinical Research Fellowship (FS/18/23/33512) and the National Institute for Health Research Oxford Biomedical Research Centre. This study was funded/supported by the NIHR Biomedical Research Centre at University Hospitals Bristol NHS Foundation Trust and the University of Bristol (GDS and TRG). The views expressed in this publication are those of the author(s) and not necessarily those of the NHS, the National Institute for Health Research or the Department of Health and Social Care. This work was supported by the Elizabeth Blackwell Institute for Health Research, University of Bristol and the Medical Research Council Proximity to Discovery Award. PE is supported by CRUK [C18281/A19169]. SL is funded by the Bau Tsu Zung Bau Kwan Yeun Hing Research and Clinical Fellowship (*200008682.920006.20006.400.01) from the University of Hong Kong.

We gratefully acknowledge all studies and databases that have made their GWAS summary data available for this study: arcOGEN (Arthritis Research UK Osteoarthritis Genetics), BCAC (the Breast Cancer Association Consortium), C4D (Coronary Artery Disease Genetics Consortium), CARDIoGRAM (Coronary ARtery DIsease Genome wide Replication and Meta-analysis), CKDGen (Chronic Kidney Disease Genetics consortium), DIAGRAM (DIAbetes Genetics Replication And Meta-analysis), EAGLE (EArly Genetics and Lifecourse Epidemiology Consortium), EAGLE Eczema (EArly Genetics and Lifecourse Epidemiology Eczema Consortium), EGG (Early Growth Genetics Consortium), ENIGMA (Enhancing Neuro Imaging Genetics through Meta Analysis), GCAN (Genetic Consortium for Anorexia Nervosa), GEFOS (GEnetic Factors for OSteoporosis Consortium), GIANT (Genetic Investigation of ANthropometric Traits), GIS (Genetics of Iron Status consortium), GLGC (Global Lipids Genetics Consortium), GliomaScan (cohort-based genome-wide association study of glioma), GPC (Genetics of Personality Consortium), GUGC (Global Urate and Gout consortium), HaemGen (haemotological and platelet traits genetics consortium), IGAP (International Genomics of Alzheimer’s Project), IIBDGC (International Inflammatory Bowel Disease Genetics Consortium), ILCCO (International Lung Cancer Consortium), IMSGC (International Multiple Sclerosis Genetic Consortium), ISGC (International Stroke Genetics Consortium), MAGIC (Meta-Analyses of Glucose and Insulin-related traits Consortium), MDACC (MD Anderson Cancer Center), MESA (Multi-Ethnic Study of Atherosclerosis), Neale’s lab (a team of researchers from Dr Benjamin Neale’s group, who made the UK Biobank GWAS summary statistics publically available), OCAC (Ovarian Cancer Association Consortium), IPSCSG (the International PSC study group), NHGRI-EBI GWAS catalog (National Human Genome Research Institute and European Bioinformatics Institute Catalog of published genome-wide association studies), PanScan (Pancreatic Cancer Cohort Consortium), PGC (Psychiatric Genomics Consortium), Project MinE consortium, ReproGen (Reproductive ageing Genetics consortium), SSGAC (Social Science Genetics Association Consortium), TAG (Tobacco and Genetics Consortium), TRICL (Transdisciplinary Research in Cancer of the Lung consortium) and UK Biobank.

JZ acknowledges his grandmother ChenZhu for all her support, may she rest in peace.

## Online methods

### Instrument selection

pQTLs from five GWAS (Sun *et al*., Emilsson *et al*., Suhre *et al*., Folkersen *et al*. and Yao *et al*.) ^9 15 16 17 18^ were used for the instrument selection (**Figure 1**). We first mapped SNPs to genome build GRCh37.p13 coordinates and then used the following criteria to select instruments:

- We selected SNPs that were associated with any protein (using a P value threshold ≤5×10^−8^) in at least one of the five studies, including both cis and trans pQTLs.
- Due to the complex LD structure of SNPs within the human Major Histocompatibility Complex (MHC) region, we removed SNPs and proteins coded for by genes within the MHC region (chr6: from 26Mb to 34Mb).
- We then conducted linkage disequilibrium (LD) clumping for the instruments with the TwoSampleMR R package ^29^ to identify independent pQTLs for each protein. We used r^2^ < 0.001 as the threshold to exclude dependent pQTLs in the cis (or trans) gene region.

After instrument selection, 2113 instruments were kept for further instrument validation (**Supplementary Table 1**). The instrument selection process, and the number of instruments for proteins at each step in the process, is illustrated in **Figure 1**.

We incorporated conditionally distinct signals from protein association data through systematic conditional analysis. Of the 5 studies, Sun *et al*. reported conditionally distinct results for both cis and trans pQTLs, which have been used in our study. Folkersen *et al*. have shared summary statistics, with which we performed approximate conditional analyses ourselves using GCTA-COJO ^35^, with genotype data from mothers in the Avon Longitudinal Study of Parents and Children (ALSPAC) as the LD reference panel ^69 70^ (a description of the ALSPAC cohort can be found in **Supplementary Note 6**). Conditionally independent signals in the cis region for Sun and Folkersen *et al*. were reported in **Supplementary Table 5**. The other studies have no summary statistics available or insufficient SNP density for conditional analysis.

### Instrument validation

For the 2113 instruments, we further classified them into three groups (noted as tier 1, tier 2 and tier 3 instruments) using two major instrument filtering steps: a pleiotropy test and a consistency test. More details of instrument validation, including harmonization of proteins and instruments and statistical tests for consistency can be found in **Supplementary Note 5**.

#### Test estimating instrument specificity

Absence of horizontal pleiotropy is one of the core assumptions for MR. This assumes that the genetic variant should only be related to the outcome of interest through the instrumented exposure. We noted that some SNPs were associated with more than one protein. For example, *APOE* SNP rs7412 is associated with a set of proteins such as ADAM11, APBB2 and APOB. We plotted a histogram of the number of proteins each instrument was associated with (**Supplementary Figure 6**) and considered instruments associated with more than 5 proteins as highly pleiotropic and assigned them as <u>Tier 3</u> instruments (which were excluded from all analyses). For instruments associated with fewer than (or equal to) 5 proteins, we reported the number of proteins each of them (and their proxies with LD r^2^>0.5) was associated with to indicate the level of potential pleiotropy. To further distinguish vertical and horizontal pleiotropy for these instruments, we used biological pathway information from Reactome (https://reactome.org/) and protein-protein interaction information from STRING DB (https://string-db.org/) implemented in EpiGraphDB (www.epigraphdb.org).

#### Distinguishing vertical and horizontal pleiotropic instruments using biological pathway data

Non-specific instruments may exhibit vertical pleiotropy (pQTL associated with proteins on the same pathway) or horizontal pleiotropy (pQTL associated with proteins on different pathways). Vertical pleiotropy does not violate the “exclusion restriction criterion” of MR but horizontal pleiotropy does ^22 27^. For any instrument associated with multiple proteins, if these proteins are mapped to the same biological pathway and/or a protein-protein interaction (PPI) exists between them, then, by definition, the instrument is more likely to act through vertical pleiotropy and it is more likely to be a valid instrument for MR. Consequently, as an approach to distinguish vertical from horizontal pleiotropy, we checked the number of pathways and PPIs each protein is involved in for all the instruments associated with 2 to 5 proteins. We used EpiGraphDB (http://www.epigraphdb.org) to extract the most specific (lowest level) pathway information related to each protein from Reactome ^71 72^ and high confidence PPIs from StringDB (confidence score >0.7) ^73 74^. First, we systematically evaluated the number of pathways each protein is involved in (either directly or as part of a complex), and how many PPIs they have. Note, that although the original databases are curated, we may expect some missing information. We further evaluated how many pathways and PPIs are shared between groups of proteins that are associated with the same SNP or SNPs in strong LD (r^2^>0.8). The number of shared components for each group of proteins is presented in **Supplementary Table 1**, and **Supplementary Data 2** depicts a detailed comparison within each group using Venn diagrams.

After this analysis, 68 instruments associated with multiple proteins were mapped to the same pathway (or same PPI), and were considered as valid instruments. Given there are other pathways and PPIs that may be not included in Reactome and STRING, we kept tier 1 and 2 instruments associated with 1 to 5 proteins for the main MR analysis, but we recorded the number of proteins and number of pathways these instruments are associated with as an indication of potential pleiotropy.

#### Consistency test estimating instrument heterogeneity across studies

We pooled pQTLs from 5 studies, which have employed different proteomics arrays. The two main assays were the SOMAscan aptamer-based multiplex protein array ^75^ and the OLINK ProSeek CVD array ^76^. The SOMAscan platform is based on the technology called Slow Off-rate Modified Aptamer (SOMAmer), for which reagents consist of a short single-stranded DNA sequence that incorporates a series of modifications that give the SOMAmer “protein-like” appendages. The OLINK ProSeek method is based on the highly sensitive and specific proximity extension array, which involves the binding of distinct polyclonal oligonucleotide-labelled antibodies to the target protein followed by quantification by real-time quantitative PCR. We noted some examples where SNPs were reported to be associated with a protein in one study but did not reach the genome-wide p-value threshold for statistical significance in other studies including the same protein. In these instances, we investigated whether this reflected no statistical evidence of association (in which case, this inconsistency may indicate potentially artefactual associations) or simply fluctuation of association strength with directionally consistent signals in both studies (which would provide supporting evidence for an instrument). Among the 2113 pQTLs selected as instruments, we looked up available protein GWAS results (Sun *et al*., Suhre *et al*. and Folkersen *et al*. with full GWAS summary statistics; Yao *et al* and Emilsson *et al* with pQTLs only) and found 1062 pQTLs (or proxies with r2>0.8) with association information in at least two studies (**Supplementary Table 15**). We then tested the beta-beta correlation using the Pearson correlation function in R. The results of the beta-beta correlations of SNP effects for each pair of studies and the number of SNPs included in each correlation analysis can be found in **Supplementary Table 2**. More details of the consistency test can be found in **Supplementary Note 5**.

We performed two consistency tests on the instruments which were present across studies. The first consistency test was a *heterogeneity test* using a pair-wise Z statistic to investigate whether there was statistical evidence of heterogeneity between effect sizes in different studies (for all pQTL studies included in our analysis where: 1) effect sizes were always in SD unit; 2) using similar sets of covariates). If the Z score was greater than 5 (equal to a P value of 0.001), we considered the instrument to have strong evidence of heterogeneity indicating inconsistency of effect sizes between studies. The second consistency test was a *colocalization analysis*, which estimates the posterior probability (PP) of the same protein measured in different studies sharing the same causal pQTL within a 1Mb window around the pQTL with the smallest P value. The default priors for colocalization analysis were used here (the prior probability a SNP is associated with the protein is 1×10^−4^; the prior probability a SNP is associated with the human phenotype is 1×10^−4^; and the prior probability a SNP is associated with both the protein and the phenotype is 1×10^−5^). We also applied the pair-wise conditional and colocalization analysis (PWCoCo) for regions with multiple pQTLs to avoid the assumptions of traditional colocalization approaches of just a single association signal per region (details in **Online methods**: *Pair-wise conditional and colocalization analysis*). A lack of evidence (i.e. PP<⍰80%) in the conventional colocalization and PWCoCo analysis would suggest that the pQTL reported in the two studies did not share the same causal signals within the region, therefore are not consistent between the studies. The colocalization analysis was conducted using the “coloc” R package ^34^. For instruments with SNP association information in both Sun *et al* and Folkersen *et al*, we were able to conduct colocalization analysis. However, due to lack of sufficient SNP coverage, it was not possible to conduct colocalization analysis to compare the pQTLs from the Emilsson *et al*, Suhre *et al* and Yao *et al* studies. We therefore conducted a LD check for these pQTLs instead. For proteins measured in multiple studies, we estimated the LD between the sentinel variant for each pQTL from one study and the top 30 associated SNPs of the other study in the same region. For pQTLs that showed only weak LD (r^2^ < 0.8) with any of the top 30 associated SNPs in the other study, we considered the pQTLs to not share the same causal SNP in the region and therefore be inconsistent instruments.

Instruments showing evidence of high heterogeneity across studies using either the pair-wise Z test (pair-wise Z > 5) or colocalization analysis (PP<80%), were flagged as <u>Tier 2</u> instruments.

Recognising that lack of replication and effect heterogeneity does not preclude at least one of these effects being genuine, we used these instruments separately for the follow-up genetic analyses (**Supplementary Table 3**) and reported the findings with caution. We designated instruments passing both pleiotropy and consistency tests as <u>Tier 1</u> instruments and used them as primary instruments for the MR analysis.

#### Identifying cis and trans instruments

We further split tier 1 instruments into two groups: 1) *cis-acting* pQTLs within a 500Kb window from each side of the leading pQTL of the protein were used for the initial MR analysis (defined as the cis-only analysis) ^22^; (2) *trans-acting pQTLs* outside the 500Kb window of the leading pQTL were designated as trans instruments. Whilst trans instruments may be more prone to pleiotropy, their inclusion could increase statistical power as well as the scope of downstream sensitivity analyses (e.g. tests for heterogeneity between instruments). Therefore, for the proteins with cis instruments, we also looked for additional trans instruments and if these were available, we conducted further MR analyses using both sets of instruments (defined as the “cis + trans” analysis).

For cis instruments from Sun *et al*. ^9^ and Folkersen *et al*. ^16^, we searched the original GWAS paper and found multiple conditional cis instruments using the following selection criteria:

1. The proteins have cis-acting pQTLs.
2. The cis-acting pQTLs passed our instrument selection procedure (defined as either tier 1 or 2 instruments).
3. The conditionally independent signals were reported in Sun *et al* ^9^.
4. LD clumping was conducted to remove pQTLs with high LD (r^2^<0.6).

After selection, 381 conditionally independent cis pQTLs associated with 153 proteins were selected to conduct the “multiple cis” MR analysis.

For cis instruments, we looked up their predicted consequence via Variant Effect Predictor ^77^ (VEP: https://www.ensembl.org/info/docs/tools/vep/index.html) hosted by Ensembl. We identified coding variants (including missense, stop-lost/gained, start-lost/gained and splice-altering variants), as epitope-binding artefacts driven by coding variants may yield artefactual cis pQTLs ^39^. We then conducted a sensitivity MR analysis that excluded cis instruments which are in the coding region to further avoid the potential issue of epitope-binding artefacts driven by coding variants.

### Phenotype selection

We obtained effect estimates for the association of the pQTLs with complex human phenotypes using GWAS summary statistics which were included in the MR-Base database (http://www.mrbase.org). We used the following inclusion criteria to select complex phenotypes to be analysed:

- The GWAS with the greatest expected statistical power (e.g. largest sample size / number of cases) when multiple GWAS records of the same phenotype / risk factor were available in MR-Base.
- GWAS with betas, standard errors and effect alleles for all tested variants (i.e. full GWAS summary statistics available).

Diseases were defined as primary outcomes. Risk factors were defined as secondary outcomes. After selection, 153 diseases and 72 risk factors (such as lipids and glucose phenotypes) were included as outcomes for the MR analyses (**Supplementary Table 6**).

### Causal inference and sensitivity analyses

The following sections describe the two-sample MR analyses using single or small numbers of instruments on 153 diseases and 72 risk factors (**Supplementary Table 6**). Positive associations between genetic instruments and phenotypes may indicate a number of potential scenarios: 1) the protein has a causal effect on the phenotype (the scenario of causality we wish to identify), 2) that the phenotype has a causal effect on protein (the reverse causality scenario), 3) confounding through LD between pQTLs and variants associated with the phenotype (for simplicity we refer to this as the ‘linkage disequilibrium scenario’) or 4) that the pQTL shares causal variants with the phenotype, but the association of the pQTL with the phenotype is not mediated by the hypothesised protein target (the ‘horizontal pleiotropy” scenario) (see **Supplementary Figure 3**). Most of the current sensitivity analysis methods, such as MR Egger regression ^55^ and Weighted Median ^42^, need a large number of independent instrumental SNPs in order to test for pleiotropy. Due to the small number of independent pQTLs available per protein we were therefore unable to implement these sensitivity analyses. To identify possible violations of assumptions of MR and to distinguish between the aforementioned scenarios, we therefore conducted the following sensitivity analyses: colocalization analysis ^34^, tests for heterogeneity between instrumental SNPs ^33^, bi-directional MR ^30^ and Steiger filtering ^31 32^ (**Figure 1**).

#### Estimating the causal effects of proteins on human phenotypes using MR

In the initial MR analysis, proteins were treated as the exposures and 225 complex human phenotypes as the outcomes (**Figure 1** – Estimate putative causal relationship). Due to high correlation amongst some of the tested phenotypes (e.g. coronary heart disease (CHD) and myocardial infarction), we used the PhenoSpD method ^79 80^ to provide a more appropriate estimate of the number of independent tests. We selected a p-value threshold of 0.05, corrected for the number of independent tests, as our threshold for prioritising MR results for follow up analyses (number of tests= 142,857; P< 3.5×10^−7^).

##### MR analysis using single locus instruments

Firstly, the strongest cis pQTL variants for each protein were used as the instrumental variable (described as ‘single cis’ analysis). The Wald ratio ^81^ method was used to obtain MR effect estimates. In this analysis, the MR effect estimates were sensitive to the particular choice of pQTLs, since only the most strongly associated SNPs within each genomic region were used as instruments. Burgess *et al* recently suggested that more precise causal estimates can be obtained using multiple genetic variants from a single gene region, even if the variants are correlated ^37 36^. Sun *et al* reported proteins with multiple cis instruments^9^, so after quality checking and LD clumping (r^2^<0.6), we used the remaining cis SNPs against all 225 phenotypes to further evaluate the MR findings from our initial MR analysis and identify potential novel associations (described as ‘multiple cis’ analysis) (**Supplementary Table 5**). A generalised inverse variance weighted (IVW) model considering the LD pattern between the multiple cis SNPs was used to estimate the MR effects. In this analysis, weights for the contribution of each SNP were obtained using pairwise LD (r^2^) calculations obtained from the 1000 Genomes European ancestry reference samples.

##### MR analysis using multi-locus instruments

Among the measured proteins reported in Sun *et al*, 34% had both cis and trans pQTLs and 30% had only trans pQTLs. Trans pQTLs that overlap phenotype association loci can provide information about previously unsuspected candidate proteins ^9^. Also, using both cis and trans instruments can provide additional accuracy and statistical power to detect causal effects ^82^. Therefore, as well as MR using only cis pQTLs, we also conducted MR on proteins with both cis and trans pQTLs (noted as the cis + trans MR analysis) and proteins with only trans pQTLs (noted as trans-only analysis). In the cis + trans MR analysis, we tested the protein-phenotype associations of 66 proteins with both cis and trans instruments. The IVW method was used to obtain MR effect estimates. In the trans-only MR analysis, we used 351 trans instruments for 298 proteins. The IVW method was used when two or more trans instruments were included in the analysis, whereas the Wald ratio method was used when only one trans instrument was included in the analysis.

##### MR analysis software

The majority of MR analyses (including Wald ratio, IVW, single SNP MR, bi-directional MR, MR Steiger filtering and heterogeneity test across multiple instruments) were conducted using the MR-Base TwoSampleMR R package (github.com/MRCIEU/TwoSampleMR) ^29^. The IVW analysis considering LD pattern was conducted using the MendelianRandomization R package (https://cran.r-project.org/web/packages/MendelianRandomization/index.html) ^83^. The MR results were plotted as forest plots and Miami plots using code derived from the ggplot2 package in R (https://cran.r-project.org/web/packages/ggplot2/index.html).

#### Distinguishing causal effects from genomic confounding due to linkage disequilibrium

Results that survived the multiple testing threshold in the MR analysis were evaluated using a stringent Bayesian model (colocalization analysis) to estimate the posterior probability (PP) of each genomic locus containing a single variant affecting both the protein and the phenotype ^34^ (**Figure 1** – Distinguishing causal effects from confounding due to LD). The default priors were used for the analysis. A PP > 80% in this analysis would suggest that the two association signals are likely to colocalize within the test region. Colocalization analysis is commonly conducted for cis QTLs ^7 8^ but under studied for trans QTLs. Given trans pQTLs show stronger pleiotropic effect than cis pQTLs (**Supplementary Figure 6B**) and may influence human phenotypes indirectly, this analysis is even more meaningful for pQTLs in the trans regions. We therefore applied colocalization to both cis and trans pQTLs. For protein and phenotype GWAS lacking sufficient SNP coverage or missing key information (e.g. allele frequency or effect size) in the test region, we conducted a LD check for the sentinel variant for each pQTL against the 30 strongest SNPs in the region associated with the phenotype as an approximate colocalization analysis. r^2^ of 0.8 between the sentinel pQTL variant and any of the 30 strongest SNPs associated with the phenotype was used as evidence for approximate colocalization. For all MR top findings, we treated colocalised findings (PP>=80%) as “Colocalised” and LD checked findings (r^2^>=0.8) as “LD checked”; other findings that did not pass the colocalization or LD check analysis were annotated as “Not colocalized”. For MR findings using multiple instruments (e.g. cis + trans analysis), we tested each pQTL with the phenotype separately. Only if all pQTLs colocalised with the phenotype at r^2^>=0.8 did we treat this finding as colocalised.

#### Pair-wise conditional and colocalization analysis

The presence of multiple conditionally distinct association signals within the same genomic region will influence the performance of colocalization analysis. We therefore developed an analysis pipeline to integrate conditional and colocalization approaches for regions with multiple conditionally independent pQTLs. Where there was convincing MR evidence below the P-value threshold of 3.5×10^−7^, but no good evidence of colocalization using the marginal SNP effects of the exposures and outcomes (in total 148 MR associations in both cis and trans regions), we performed pairwise colocalization analyses of all conditionally distinct pQTLs against all identified conditionally distinct association signals in the outcome data (noted as pair-wise conditional and colocalization analysis: PWCoCo). The conditional analysis for proteins and human phenotypes was conducted using the GCTA-COJO package^35^, with genotype data from mothers in the Avon Longitudinal Study of Parents and Children (ALSPAC) as the LD reference panel ^69 70^ (a description of the ALSPAC cohort can be found in **Supplementary Note 6**). **Figure 2** demonstrates the 9 possible pair-wise combinations of various conditional signals for proteins and phenotypes at which there are 2 independent signals in the region (the 9 combinations listed in the following 3×3 table).

###### Box 1. The 9 possible pair-wise combinations of various conditional signals for proteins and phenotypes in a 3×3 table

|  | Marginal effect of the exposure (M) | Joint effect of the exposure conditional on the top hit (C1) | Joint effect of the exposure conditional on second hits (C2) |
| --- | --- | --- | --- |
| Marginal effect of the outcome (M) | MvM | C1vM | C2vM |
| Joint effect of the outcome conditional on the top hit (C1) | MvC1 | C1vC1 | C2vC1 |
| Joint effect of the outcome conditional on second hits (C2) | MvC2 | C1vC2 | C2vC2 |

For protein-phenotype associations which only showed colocalization evidence after we applied PWCoCo, we recorded the PWCoCo model which showed colocalization evidence in a new column “PWCoCo_model”, in **Supplementary Tables 7, 8, 11, 12, 13, 16 and 17.**

#### Heterogeneity test of MR findings

For MR analyses using two or more instruments, we conducted heterogeneity tests to estimate the variability in the causal estimates obtained for each SNP (i.e. how consistent is the causal estimate across all SNPs used as separate instruments) (**Figure 1 —** Consistency of the causal estimate across all SNPs). Cochran’s Q test statistic was calculated for the IVW analyses, which is expected to be chi-squared distributed with number of SNPs minus one degrees of freedom ^33^. Lower heterogeneity suggests a lower chance of violations of assumptions in MR estimates, such as the presence of confounding through horizontal pleiotropy ^84^.

#### Distinguishing causal effects for proteins on phenotypes from reverse causality

With sufficiently large sample sizes, a SNP associated with an outcome through a mediating exposure could reach the conventional threshold for statistical significance in both the outcome and exposure GWAS. Therefore, using such thresholds to define instruments could lead to situations where the instrumental SNP influences the hypothesised exposure via the hypothesised outcome (i.e. the hypothesised outcome actually has a causal effect on the hypothesised exposure and not vice versa). In order to mitigate the potential impact of this limitation, we used two approaches to identify directions of causality: bi-directional MR and Steiger filtering.

##### Reverse Mendelian randomization

For associations between proteins and phenotypes identified in the MR analysis, we applied bi-directional MR to evaluate evidence for causal effects in the reverse direction by modelling complex phenotypes as our exposure and plasma protein as our outcome. Instruments for complex phenotypes were selected based on a threshold of P < 5 × 10^−8^ from GWAS after LD clumping to identify independent variants. The IVW method was applied to estimate the causal effects of phenotypes on proteins where more than one instrument was available, otherwise the Wald ratio was used. MR-Egger ^55^ was used as a sensitivity analysis to test for potential pleiotropic effects.

##### Identifying the direction of effects for instruments using Steiger filtering

Due to lack of sufficient SNP association information (e.g. allele information, effect size, standard error) for some pQTL studies, it was not possible to conduct bi-directional MR using all proteins as outcomes. Therefore, we conducted Steiger filtering as an alternative method to test the directionality of protein-phenotype associations. The Steiger method ^85^ has been implemented in the TwoSampleMR R package ^29^ to assess directionality of instrument-phenotype associations ^31 32^.

The process of choosing valid instruments using Steiger filtering follows these steps:

1. Select the top findings from all five studies using a p-value threshold of 3.5 × 10^−7^ (which is the Bonferroni P value threshold of the MR analysis).
2. Classify instruments in each MR analysis based on Steiger filtering:
  - ‘TRUE’: evidence for causality in the expected direction i.e. protein precedes phenotype.
  - ‘FALSE’: evidence for causality in the reverse direction i.e. phenotype precedes protein. Instruments with ‘FALSE’ were removed from the sensitivity analysis.
  - ‘NA’: no result (due to insufficient summary data from the study to estimate the SNP-protein and/or SNP-phenotype correlation, e.g. missing effect allele frequencies in the outcome data or missing numbers of cases and controls for binary phenotypes).

For disease phenotypes, we estimated the variance explained on the liability scale. Based on step 2, we set up a flag (categorical variable) to record the direction of the effects of the SNPs using Steiger filtering.

Steiger filtering acts slightly different for MR using cis or trans pQTLs. For cis pQTLs, measurement error may bias the results. For trans pQTLs, a confounder may bias the results. However, the bias from these issues is expected to be minimal.

#### Drug target validation and repositioning

Approved drug targets have previously been shown to be enriched for gene-phenotype associations ^6^. We therefore wished to assess whether approved drug targets were enriched for protein-phenotype associations, as obtained in the present study using MR. We assessed the support for approved drug targets among our MR findings using Fisher’s exact test. Target-indication pairs for successful and failed drugs were identified using a manually annotated version of PharmaProjects database from Citeline (https://pharmaintelligence.informa.com/). The phenotypes used in the MR analyses and the indications listed in Citeline’s PharmaProjects (downloaded on the 9th of May 2018) were then manually mapped to MeSH headings as a common ontology. This allowed us to match the protein-phenotype associations with corresponding target-indication pairs. To improve this matching, we implemented a similarity matrix, derived from all MeSH headings in the manual mapping, and retained matches with a relative similarity greater than 0.7 for our analyses (the similarity matrix has been previously described in Nelson *et al*. ^6^). We then compared whether the target-indication pair represented a successful or failed drug against whether there was a signal or not for the corresponding protein-phenotype pair among our MR findings. For the purposes of this test, a signal was defined as an MR result with a p-value less than 3.5 × 10^−7^ (which is the Bonferroni P value threshold of the MR analysis) with supporting evidence from colocalization analysis. We further conducted a set of sensitivity analyses based on the following criteria to increase the reliability of the enrichment analysis:

1. We checked the direction of effect of MR findings and drug trial results for the 8 approved drugs using therapeutic direction information from PharmaProjects.
2. For target-indication pairs linked to similar phenotypes (for example, the same target associated with angina and myocardial infarction), we removed one of them to avoid double counting the same association.
3. To avoid the influence of epitope-binding artefacts, we removed MR results estimated using missense variants as an instrument.
4. We checked whether approved drugs had been motivated by genetics from Drug Bank (https://www.drugbank.ca/), which may have inflated the OR estimate.

In total, we removed 75 target-indication pairs based on criteria 2 (45 pairs), 3 (23 pairs) and 4 (2 pairs; some pairs appeared in multiple situations) and conducted the comparison between protein-phenotype associations using MR and target-indication pairs from PharmaProjects, both on each criterion separately and on all criteria together (**Supplementary Table 21**).

Phenome-wide MR has demonstrated the potential to validate, repurpose and predict on-target side effects of drug targets. Of the protein-phenotype associations that showed evidence of colocalization identified in the cis-only, cis+trans, trans-only or MR analyses using pQTLs with heterogeneous effects across studies (noted as Tier 2 instruments), we first looked up how many proteins with MR evidence were established drug targets in the Informa PharmaProjects database. We then looked up how many of the associations were established target-indication pairs in the PharmaProjects database. More importantly, we predicted the potential adverse effects and repositioning opportunities of all marketed drugs and drugs under development using phenome-wide MR. The forest plots illustrating phenome-wide MR results were drawn using the R package “ggplot2” (https://ggplot2.tidyverse.org/).

#### Enrichment of proteome-wide MR with the druggable genome

Previously, Finan *et al* systematically identified 4479 genes as the newest druggable genome compendium ^46^. This study stratified the druggable genome set into three tiers. Tier 1 (1427 genes) included efficacy targets of approved small molecules and biotherapeutic drugs, as well as targets modulated by clinical-phase drug candidates; tier 2 was composed of 682 genes encoding proteins closely related to drug targets, or with associated drug-like compounds; and tier 3 contained 2370 genes encoding secreted or extracellular proteins, distantly related proteins to approved drug targets, and members of key druggable gene families not already included in tier 1 or tier 2. We assessed whether the 1002 proteins we selected for the MR analyses overlapped with the 4479 genes from the druggable genome (**Supplementary Table 24**). The proteins were mapped based on the HGNC name of the encoding genes. We further assessed the overlap based on whether the protein had cis or trans instruments and based on the druggable genome tiers.

In addition to the above comparison between instrumentable and druggable genome, we also assessed the enrichment of top pQTL MR findings with the druggable genome. 295 protein-phenotype associations (120 proteins on 70 phenotypes) with both MR and colocalization evidence were selected for this analysis. We stratified the 120 proteins into 4 groups based on their druggability: tier 1 contained 23 proteins, tier 2 contained 11 proteins, tier 3 contained 58 proteins, and 28 proteins remained unclassified. The 70 phenotypes were stratified into 8 groups: 8 autoimmune diseases, 3 bone phenotypes, 8 cancer phenotypes, 12 cardiovascular phenotypes, 4 glycemic phenotypes, 2 lung phenotypes, 4 psychiatric phenotypes and 29 other phenotypes. The protein-phenotype associations with MR and colocalization evidence were coloured separately based on their druggability tiers. More details of this enrichment analysis are shown in **Supplementary Table 25 and Figure 5**.

## Data availability

The data (GWAS summary statistics) used in the analyses described here are freely accessible in the MR-Base platform (www.mrbase.org). All our analysis results for 989 proteins against 225 human phenotypes are freely available to browse, query and download in EpiGraphDB (http://www.epigraphdb.org/pqtl/). An application programming interface (API) and R package documented on the website enable users to programmatically access data from the database.

## Code availability

The code used in the Mendelian randomization analyses described here are freely accessible in the TwoSampleMR R package via GitHub (https://github.com/MRCIEU/TwoSampleMR). Full documentation for the R package is also provided (https://mrcieu.github.io/TwoSampleMR/). We implemented the colocalization analysis using the coloc R package (created by Chris Wallace *et al*.), which can be downloaded here (https://cran.r-project.org/web/packages/coloc/index.html).

