## Supplementary documents for "Phenome-wide Mendelian randomization mapping the influence of the plasma proteome on complex diseases"

### Supplementary Tables

All supplementary tables can be found in the file “pQTL-MR-Supplementary-tables”. The colours of header line for all tables refer to different type of information. Protein (exposure) information in pink; instrument information in yellow; SNP association information in green; Mendelian randomization results in blue; follow-up and sensitivity analysis information in grey (e.g. colocalization and heterogeneity analysis results); outcome information of human traits in purple; trial information in red.

**Table legends**

Supplementary Table 1. Genetic instruments for plasma proteins which were included in the validation process. Each instrument contains four types of information: protein/exposure information, instrument information, SNP association information and instrument validation information.

Supplementary Table 2. The pair-wise correlations of SNP effects across pQTL studies. For the 1062 SNPs with SNP effects in two studies, a pair-wise comparison using Pearson correlation (r) were conducted. The overall agreement of SNP effects across each pair of studies was estimated.

Supplementary Table 3. Tier 2 genetic instruments for plasma proteins. Tier 2 instruments are defined as SNPs which passed the specificity test but show heterogeneity across different studies. Each instrument contains four types of information: protein/exposure information, instrument information, SNP association information and instrument validation information.

Supplementary Table 4. Cis and trans genetic instruments for plasma proteins. Tier 1 instruments that have both cis and trans acting SNPs for the same protein, which were used in the multiple instrument analysis. Each instrument contains four types of information: protein/exposure information, instrument information, SNP association information and instrument validation information.

Supplementary Table 5. Multi-cis genetic instruments for plasma proteins. Tier 1 instruments with more than one cis associated SNP for the same protein, which were used for the multiple instrument MR analysis. Each instrument contains four types of information: protein information, instrument information, SNP association information and instrument validation information.

Supplementary Table 6. Diseases and risk factors used as outcomes in our study. The 225 phenotypes were unique phenotypes selected from MR-Base. For phenotypes with multiple studies in MR-Base, we selected the GWAS with the largest sample size if the outcome was continuous or the GWAS with the largest number of cases if the outcome was discrete (disease outcomes).

Supplementary Table 7. The top findings for cis-only Tier 1 instruments. These associations showed evidence of putative causal relationship between proteins (with cis and Tier 1 instruments) and human traits in both the Mendelian randomization (P<3.5x10^-7^) and colocalization analysis (Probability of colocalization >80%). Notation: “Share_same_pathways?” refers to whether the proteins linked to the same pQTL share the same biological pathway; “N_shared_PPIs” is the number of pair-wise protein-protein interactions (PPIs) each protein is involved in; “Shared_proteins” lists the names of the interacting proteins; “N_proteins_groups” is the number of protein group each instrument is associated with after accounting for proteins in the same biological pathway and/or with PPI to each other. Steiger_filter and Steiger_P refer to whether the pQTL pass the Steiger filtering and the P value of the Steiger filtering analysis. VEP refers to the functional consequence of the pQTL obtained using Variant Effect Predictor (VEP). WR under Method column refers to the Wald ratio approach. Conditional analysis refers to whether the colocalization analysis was conducted on the original SNP effects or the conditioned SNP effects independent of the protein and phenotype. PP.H0.abf to PP.H4.abf refers to the posterior probabilities of the 5 colocalization hypotheses, H0: neither pQTL or outcome is associated in the region; H1: only pQTL is associated in the region; H2: only outcome is associated in the region; H3: both phenotypes are associated, but with different causal variants; H4: both phenotypes are associated, and share a single causal variant. Outcome_SNP and LD_r2 under LD check refer to the proxy SNP of the outcome with highest LD to the pQTL within the 1Mb window as well as the r^2^ between proxy SNP of the outcome and the pQTL. Type	under Coloc? column refers the type of colocalization evidence, “LD checked” means the colocalization evidence come from the LD check analysis where “Colocalised” means the colocalization evidence come from the colocalization analysis. Reported_effector_gene? Column refers to whether the protein coding gene have previously been reported to be associated with the outcome phenotype.

Supplementary Table 8. The top findings for cis-only Tier 1 instruments without colocalization evidence. These results showed evidence of putative causal relationship between proteins (with cis and Tier 1 instruments) and human traits using Mendelian randomization but did not show enough evidence of colocalization across the traits.

Supplementary Table 9. The top Mendelian randomisation results using multiple cis instruments. For proteins with multiple independent cis QTL this analysis implements IVW MR to generate a combined estimate. A) MR findings which showed similar association using single cis analysis and multiple cis instruments; B) MR findings which showed increased power using multiple cis instruments comparing to MR using single cis instrument. Notation: IVW_SB under Method column refers to the inverse variance weighted method accounting for the correlation between multiple cis instruments, which was implemented by the MendelianRandomization R package (developer: Olena Yavorska and Stephen Burgess)

Supplementary Table 10. The Mendelian randomization and colocalization results of Haptoglobin (HP) on LDL cholesterol (LDL-C) for both cis HP instruments.

Supplementary Table 11. The top Mendelian randomization results using both cis and trans acting instruments.

Supplementary Table 12. The top Mendelian randomization results using trans-only instruments.

Supplementary Table 13. The top Mendelian randomization results using trans-only instruments without colocalization evidence.

Supplementary Table 14. The heterogeneity test for the 47 proteins with multiple trans-pQTLs.

Supplementary Table 15. Details of the instrument validation results across studies. For all instruments identified from the 5 studies, a SNP lookup were conducted. There were 1062 SNPs (or proxies with r^2^>0.8) with available association information in one or more other studies using the protein ID mapping we conducted (shown in pink background). Protein associated SNP reported in the original paper were noted as “pQTL”. Protein associated SNP we found in additional studies were called “Lookup_SNP”. Effect allele, other allele, effect allele frequency, beta, standard error, P value and sample size of the pQTLs and Lookup SNPs were highlighted in green and blue separately. For these SNPs, two consistency analyses were conducted: 1) the heterogeneity test using a pair-wise Z test to investigate whether there was statistical evidence of heterogeneity between effect sizes in different studies (results shown in light grey background); 2) the colocalization analysis estimates the Posterior Probability (PP) of the same protein measured in different studies sharing the same causal pQTL within a 2Mb window around the pQTL (results shown in dark grey background).

Supplementary Table 16. The top Mendelian randomization results using Tier 2 instruments with evidence of heterogeneity for instruments across studies. These instruments were lack of replication and effect heterogeneity but didn’t preclude these effects being genuine. The MR analyses were conducted using the most significant SNP across studies.

Supplementary Table 17. The top Mendelian randomization results using Tier 2 instruments with not enough evidence of colocalization for same proteins across studies. These instruments did not show enough evidence of colocalization, which suggested two different signals within the test region. Therefore, when possible, the MR analysis were performed using instruments from each study separately in case one of the two distinct signals was truly associated with the outcome.

Supplementary Table 18. The eQTL and immune response QTL lookup results for two cis pQTLs of IL18R1. The eQTL and immune response QTL data were from the GTEX (sample size (n) from 80 to 491 for 48 tissues), the eQTLGen (n=31,684) and Kim-Hellmuth et al. (n=134). Since the second pQTL rs13014644 were missing in GTEX, we used a SNP with perfect LD to it as a proxy (rs1946131, r^2^=1 in the 1000 Genome Europeans). A) the “Tissue” column refers the relevant tissue type were the eQTL were measured; B) the “Condition” column refers various simulation stages of the monocytes, LPS, MDP, and RNA refer to three defined microbe-associated molecular patterns of the stimulate cells, 90m and 6h refer to different time points (90 minus and 6 hours). All eQTL records with P value < 1x10^-4^ were highlighted in red.

Supplementary Table 19. The drug target validation results. In this analysis we cross validated our top Mendelian randomization findings with trial information from PharamProjects database.

Supplementary Table 20. Approved drugs with MR and colocalization evidence. This table shown the 8 approved target-indication pairs with MR and colocalization evidence. The MR, colocalization, trial information and MeSH term mapping information of this comparison have been provided.

Supplementary Table 21. Enrichment analysis comparing drug trial evidence with Mendelian randomization and colocalization evidence. Note: The protein-trait association pairs were grouped into four categories: 1) pairs with both MR + colocalization and drug trial evidence; 2) pairs with MR + colocalization evidence but no drug trial evidence; 3) pairs with no strong MR evidence but with drug trial evidence; and 4) pairs with no MR or drug trial evidence. The cut-off for MR evidence was p< 3.5x10^-7^; the cut off for colocalization evidence was probability > 80%. The drug trial evidence was obtained from PharmaProjects database. A Fisher’s exact test were then conducted in this enrichment analysis to test whether the target-indication pair represented a successful or failed drug against a signal or not for the corresponding protein-trait pair among the MR and colocalization findings. The MR and colocalization analysis results involved in this analysis including 1) Tier 1 instruments in the cis region; 2) Tier 1 and 2 instruments in the cis region; and 3) both Tier 1 and Tier 2 instruments in both cis and trans region.

Supplementary Table 22. Potential repurposing opportunities of approved drugs and novel drugs under development. In this supplementary table, we listed protein-trait associations for novel drugs under development to additional indications. The MR, colocalization, trial information and MeSH term mapping information have been provided.

Supplementary Table 23. Drugs in development with MR and colocalization evidence. This table shows predicted efficacy of drugs under development. The MR, colocalization, trial information and MeSH term mapping information have been provided.

Supplementary Table 24. The overlap between 1002 instrumented proteins and the druggable proteins. The druggable genome information was obtained from Finan et al. The druggable proteins had been grouped into three tiers: Tier 1 (1427 genes) included efficacy targets of approved small molecules and biotherapeutic drugs as well as clinical-phase drug candidates; tier 2 was composed of 682 genes encoding targets with known bioactive drug-like small-molecule binding partners; and tier 3 contained 2370 genes encoding secreted or extracellular proteins, proteins with more distant similarity to approved drug targets, and members of key druggable gene families not already included in tier 1 or 2.

Supplementary Table 25. The overlap between protein-phenotype associations from MR and the druggable proteins. We selected 295 protein-phenotype associations (120 proteins on 70 phenotypes) with both MR and colocalization evidence for this analysis. For the 120 proteins, we grouped them into 4 groups based on the evidence level of being druggable: tier 1 contained 23 proteins, tier 2 contained 11 proteins, tier 3 contained 58 proteins, and no tier contained 28 non-druggable proteins (with current druggable evidence). The 70 phenotypes were grouped into 8 categories: 8 autoimmune diseases, 3 bone phenotypes, 8 cancer outcomes, 12 cardiovascular phenotypes, 4 glycemic phenotypes, 2 lung phenotypes, 4 psychiatric phenotypes and 29 other phenotypes. The protein-phenotype associations with MR and colocalization evidence were coloured separately based on their druggability tiers.

Supplementary Table 26. The Mendelian randomization results for MMP12 on cardiovascular diseases. The cis instrument rs28381684 associated with MMP12 were used to test the association for MMP12 on coronary heart disease and stroke (and stroke subtypes).

Supplementary Table 27. The protein mapping across four studies. The somalogic id was not reported in Emilsson et al, therefore the protein mapping information for this study could not be included.

### Supplementary Figures

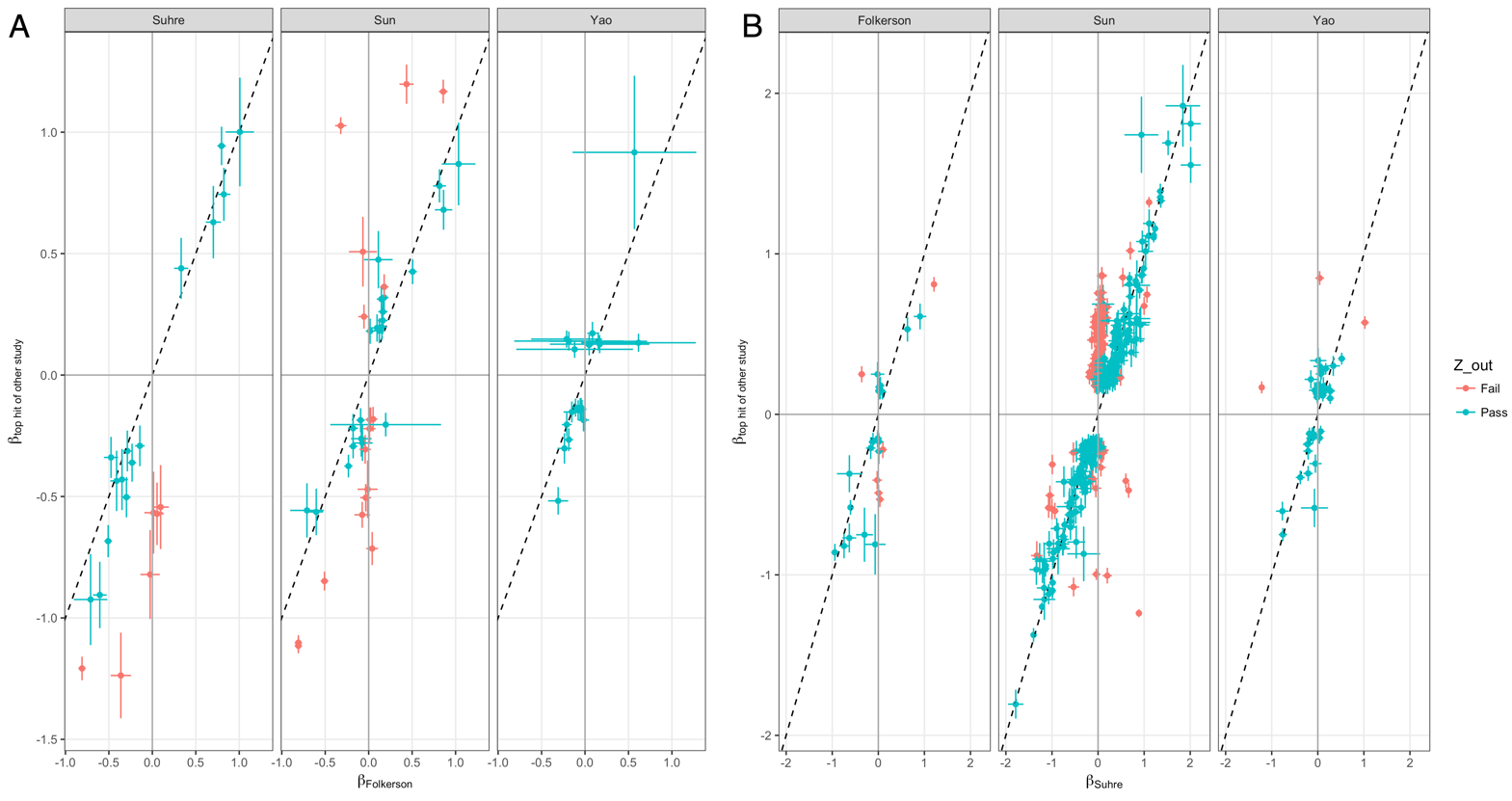

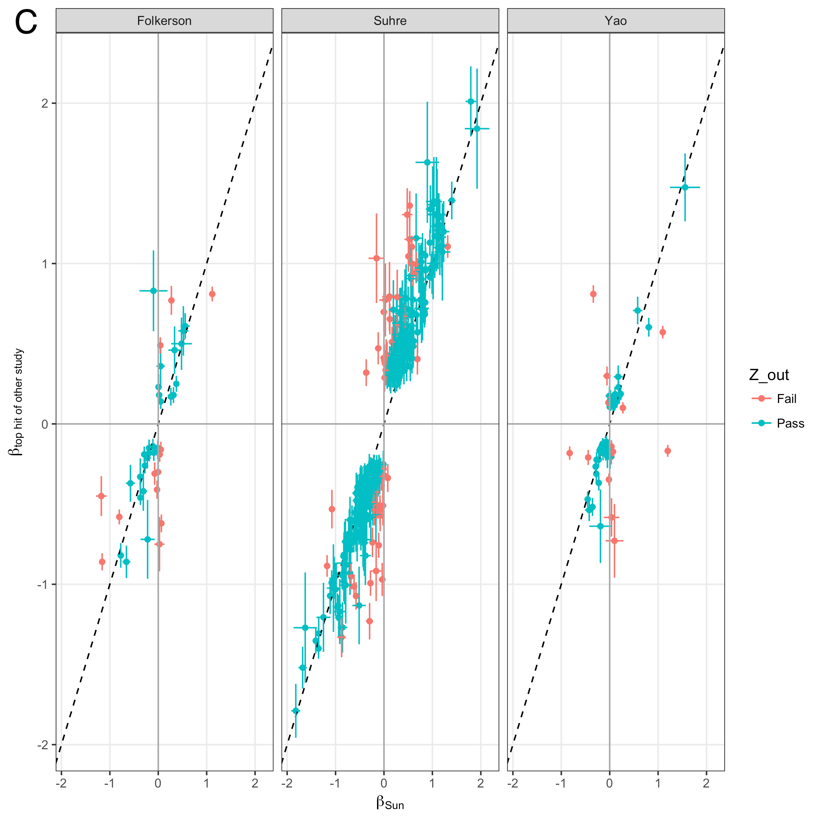

**Supplementary Figure 1.** Scatter plot showing the pair-wise instrument validation across the pQTL GWAS used in the MR study. A) SNP effects from Folkersen vs SNP effects from Sun et al, Suhre et al and Yao et al; B) SNP effects from Suhre vs SNP effects from Sun et al, Folkersen et al and Yao et al; C) SNP effects from Sun vs SNP effects from Folkersen et al, Suhre et al and Yao et al. Notation: X-axis refers to the SNP effects from the lookup study; Y-axis refers to the SNP effects from the other studies; dotted line is the identity line. Each dot refers to one SNP in the comparison, dots in green refer to the SNP effects which have a pair-wise Z score > 5, whereas the dots in red refer to SNP effects which have a pair-wise Z score < 5.

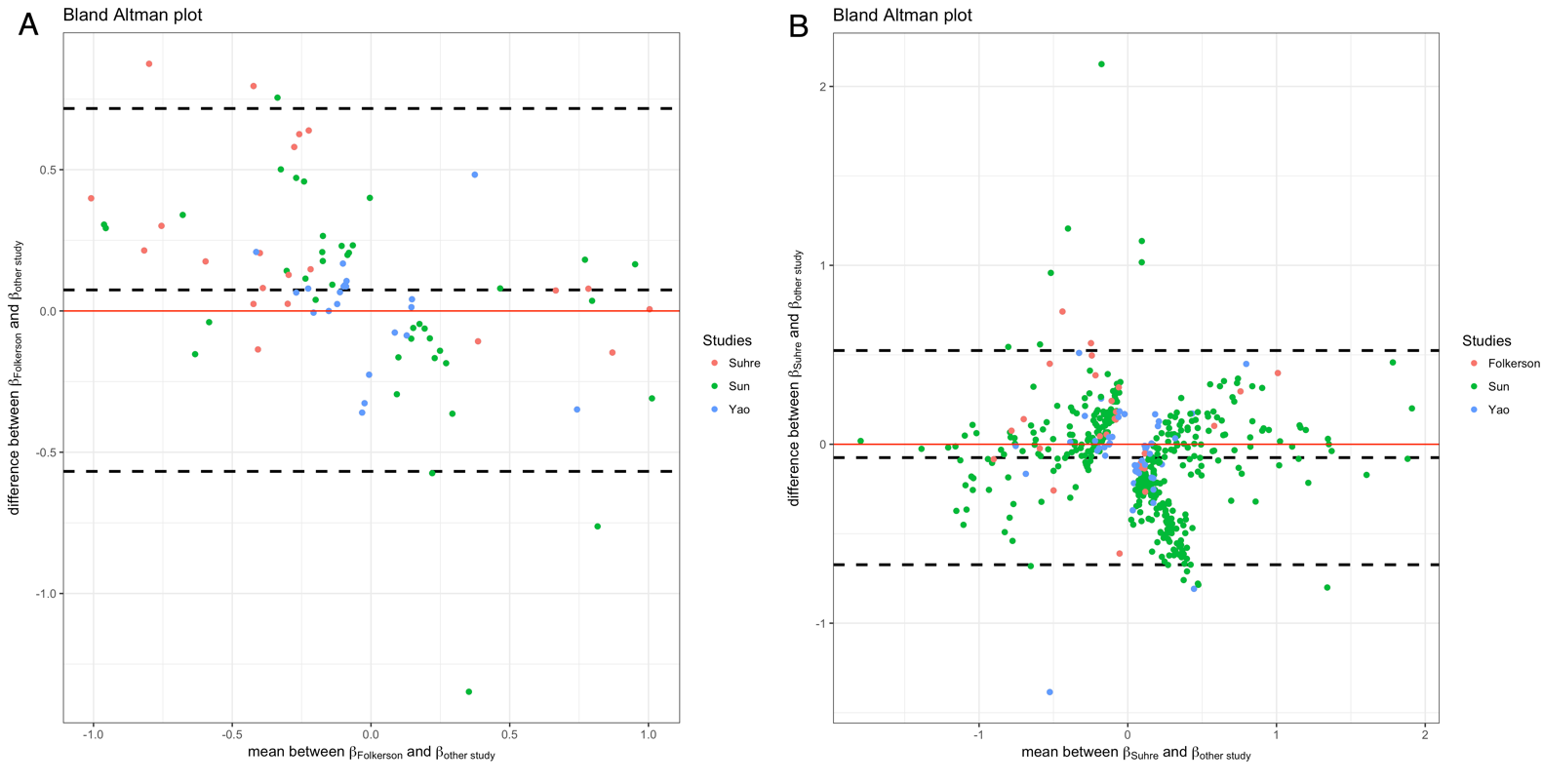

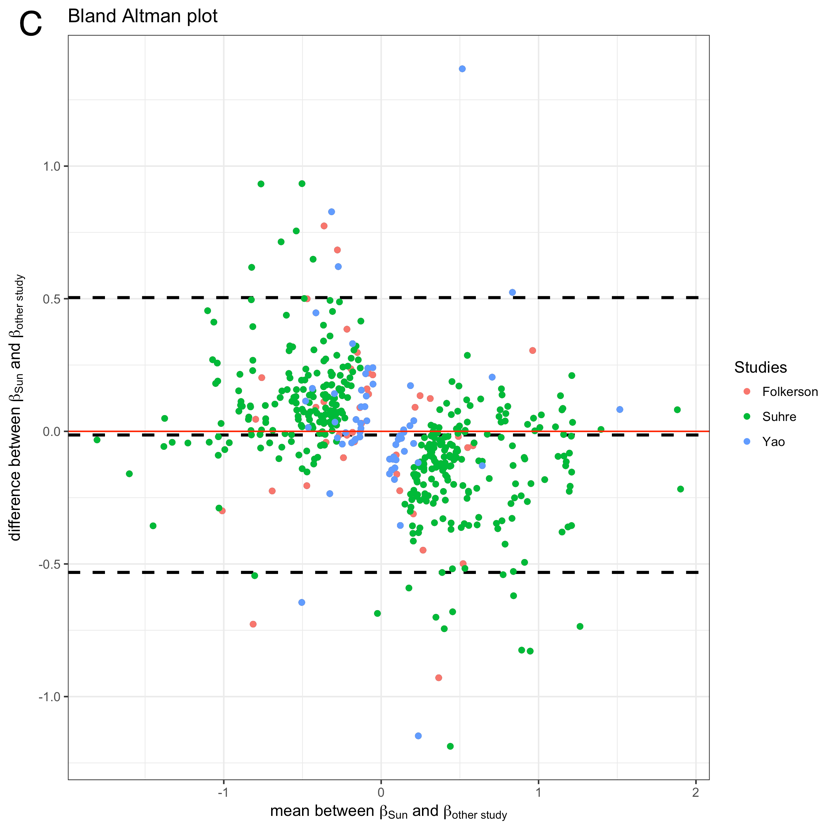

**Supplementary Figure 2.** Bland Altman plots showing pair-wise instrument validation across the pQTL GWASs used in the MR study. A) SNP effects from Folkersen vs SNP effects from the other three studies; B) SNP effects from Suhre vs SNP effects from the other three studies; C) SNP effects from Sun vs SNP effects from the other three studies. Notation: X-axis refers to the mean across effects; Y-axis refers to the difference between effects. The three dotted lines refer to the central estimate and the 95% confidence interval lines of the Bland Altman test. Each dot refers to one SNP in the comparison, dots in red, green and blue refer to the SNP effects from each of the different studies.

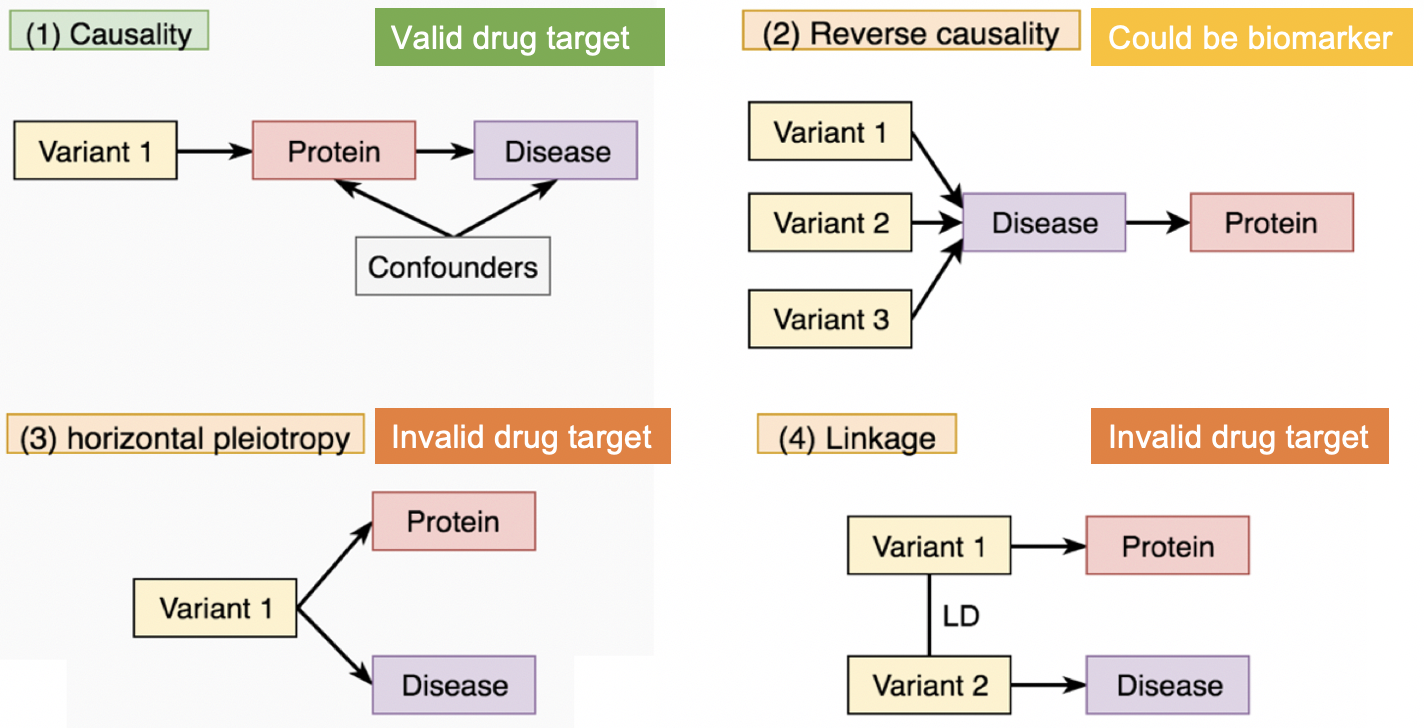

**Supplementary Figure 3.** MR and colocalization models. Model 1 – Causality: A genetic variant affects disease risk by changing protein levels; Model 2: Reverse causality Genetic variants affect disease risk through pathways other than via the protein of interest. The disease has a downstream effect on protein levels; Model 3 – Horizontal pleiotropy: a genetic variant influences both protein levels and disease risk by two independent biological pathways; Model 4 – confounding by LD: a genetic variant (variant 1) that influences protein levels is correlated with a second variant (variant 2) that influences disease risk. Colocalization analysis can distinguish Model 4 from Model 1 or Model 3.

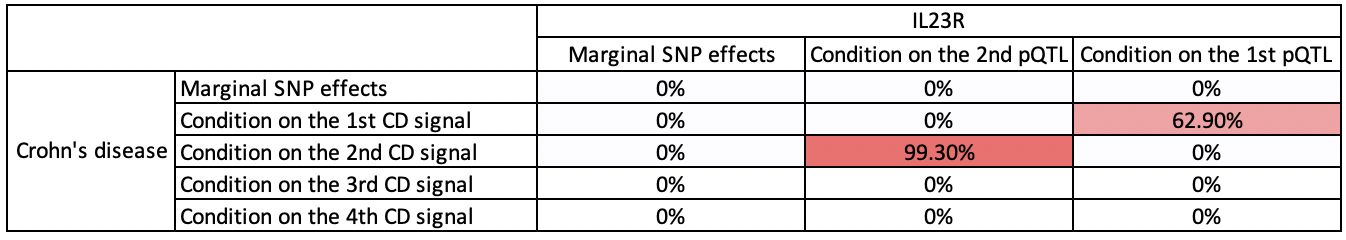

**Supplementary Figure 4.** Heatmap of the colocalization evidence for IL23R association on Crohn’s disease (CD) in the IL23R region. The 15 cells refer to the 15 pair-wise combinations of pair-wise conditional and colocalization analysis. The three columns refer to the SNP effects of IL23R protein level used in the colocalization analysis (marginal SNP effect, joint SNP effect after conditioning on the 2^nd^ IL23R signal (rs3762318) and the joint SNP effect after conditioning on the 1^st^ IL23R signal (rs11581607)). The five rows refer to the SNP effects of Crohn’s disease used in the colocalization analysis (marginal SNP effect, joint SNP effects after conditioning on the 1^st^, 2^nd^, 3^rd^ and 4^th^ CD signals). The darker red colour refers to stronger colocalization evidence.

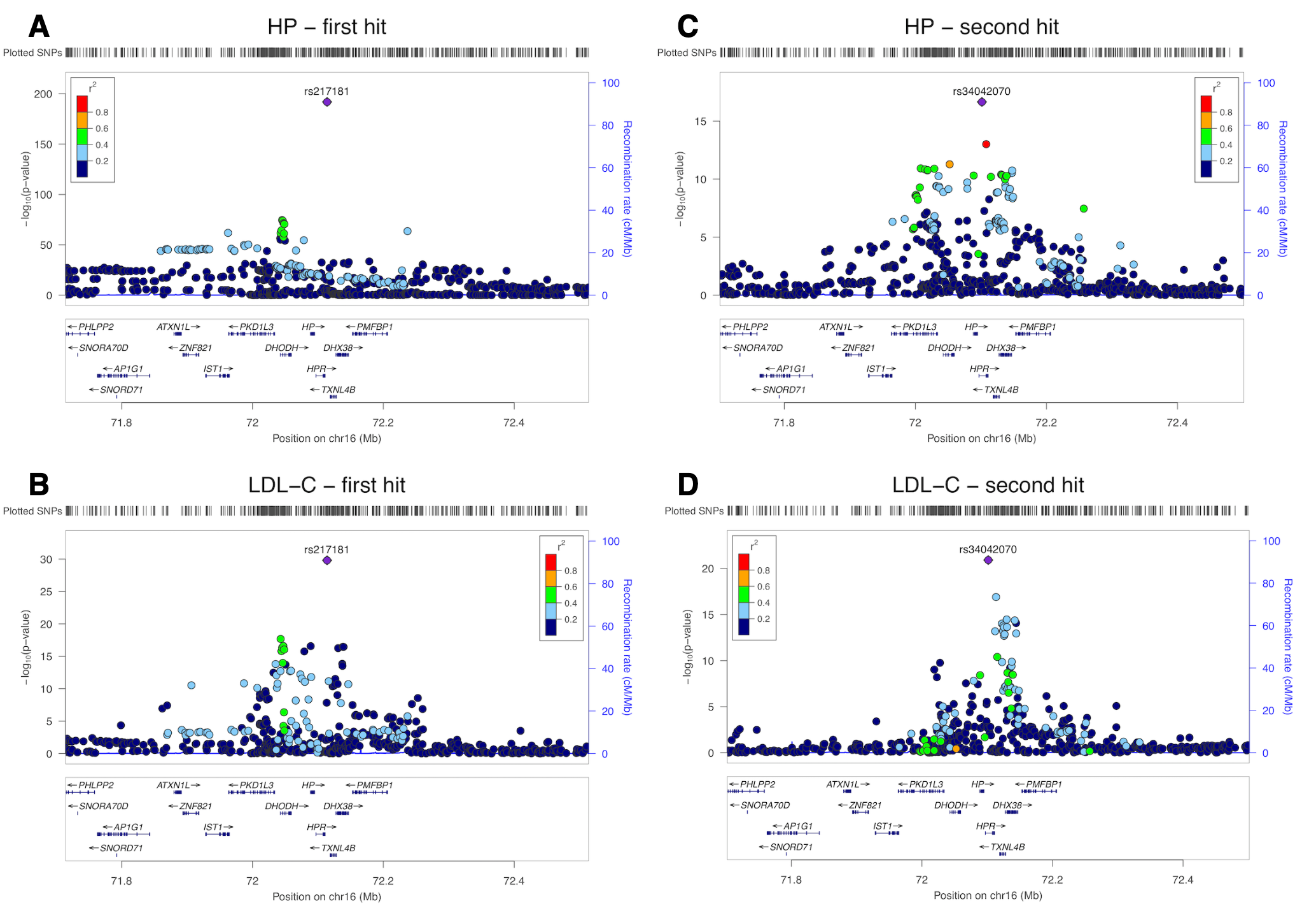

**Supplementary Figure 5.** Regional association plot showing multiple association peaks for Haptoglobin (HP) and LDL cholesterol in the cis region. The second independent hit for LDL cholesterol colocalised with the top hit for HP (rs217181) after conditioning on the top hit for LDL cholesterol (rs2000999) (colocalization probability = 99.9%). (A) HP after condition on the second pQTL HP rs34042070; (B) LDL-C after condition on the second pQTL HP rs34042070; (C) HP after condition on the top pQTL HP rs217181; (D) LDL-C after condition on the top pQTL HP rs217181.The HP data are from Sun et al. and the LDL cholesterol data are from GLGC consortium.

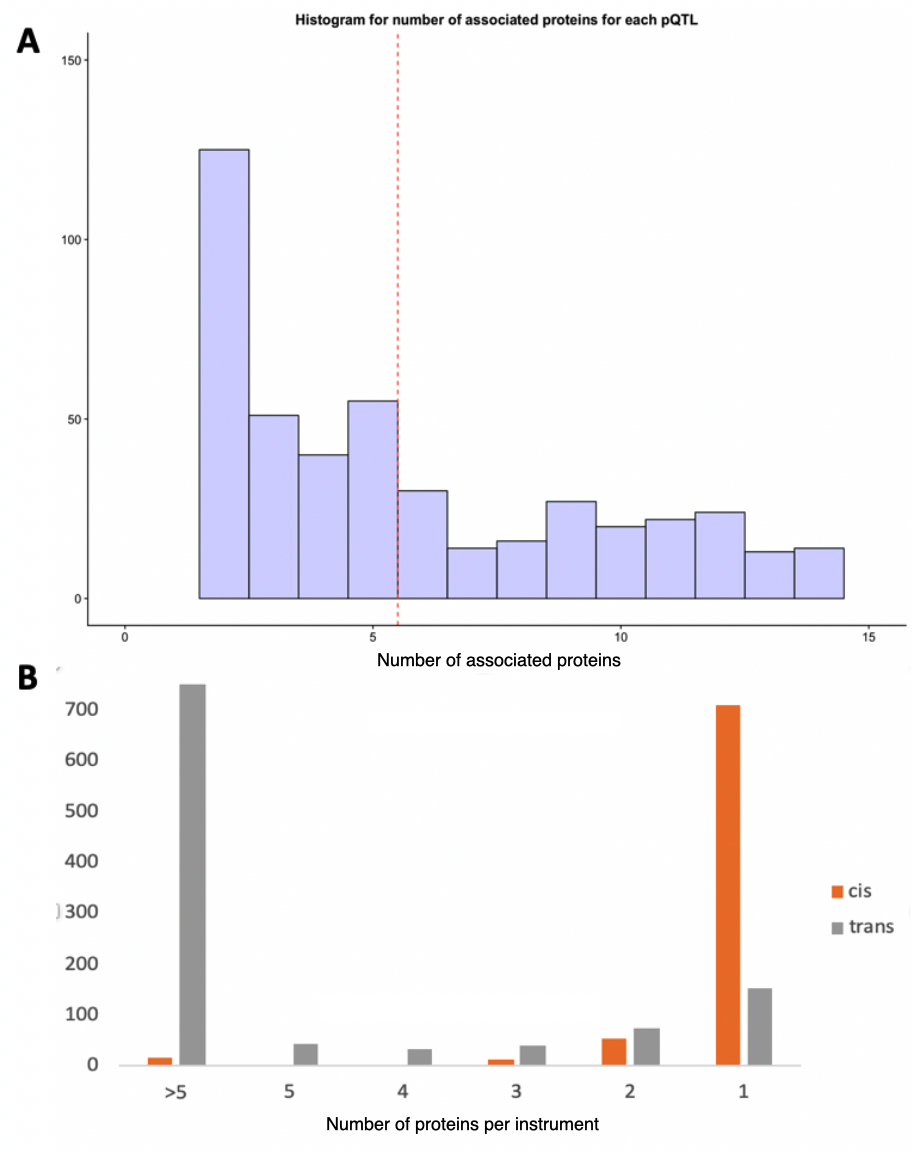

**Supplementary Figure 6.** Histogram representing the distribution of the number of proteins associated with each pQTL. (A) a zoomed in histogram which only showed pQTLs associated with fewer than 15 proteins. (B) a histogram of number of proteins each pQTL associated with, the grey bar refers to trans pQTL; the orange bar refers to cis pQTL. There is a clear trend that the trans pQTLs were associated more proteins (potentially pleiotropic) compared to the cis pQTLs.

A

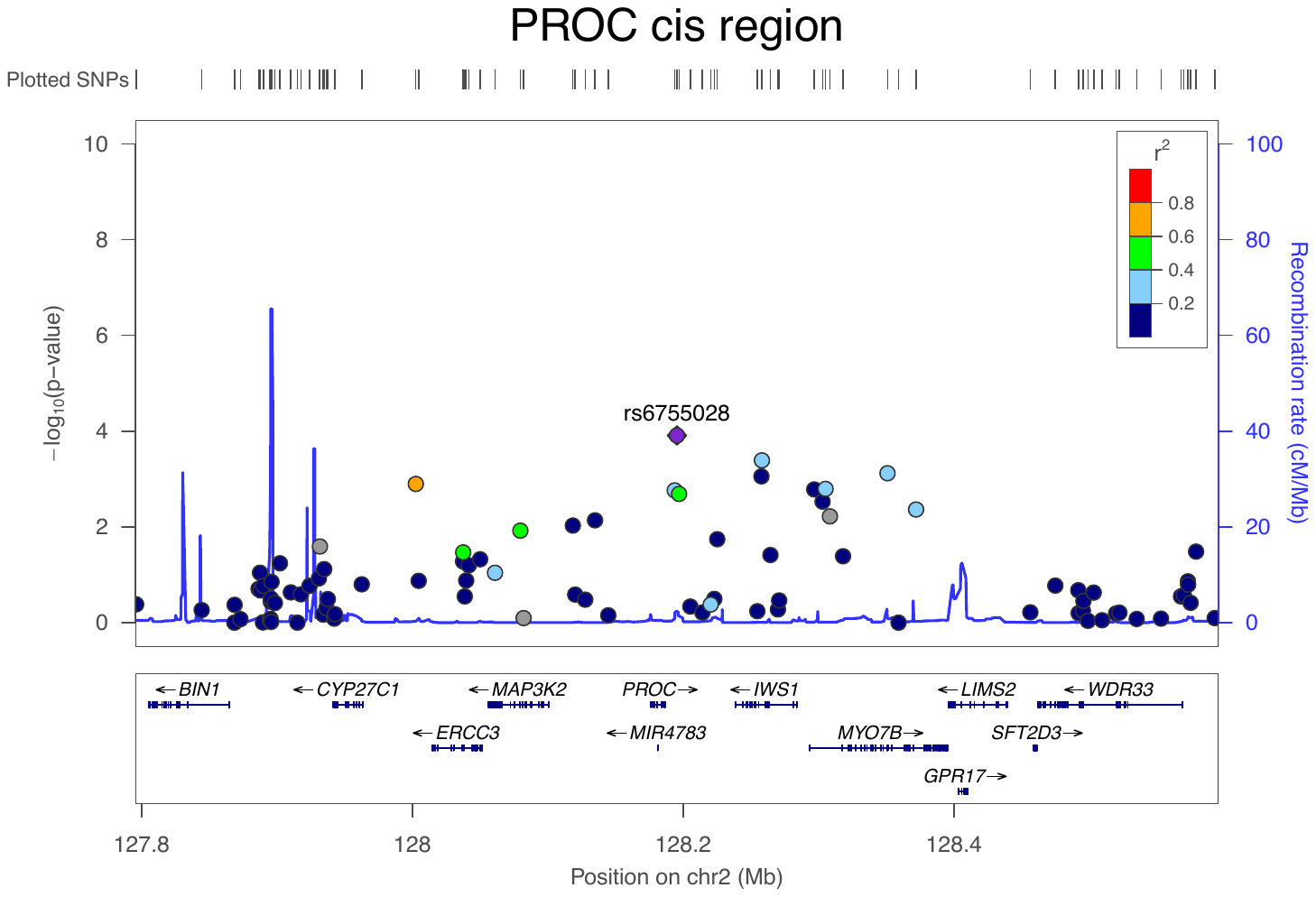

B

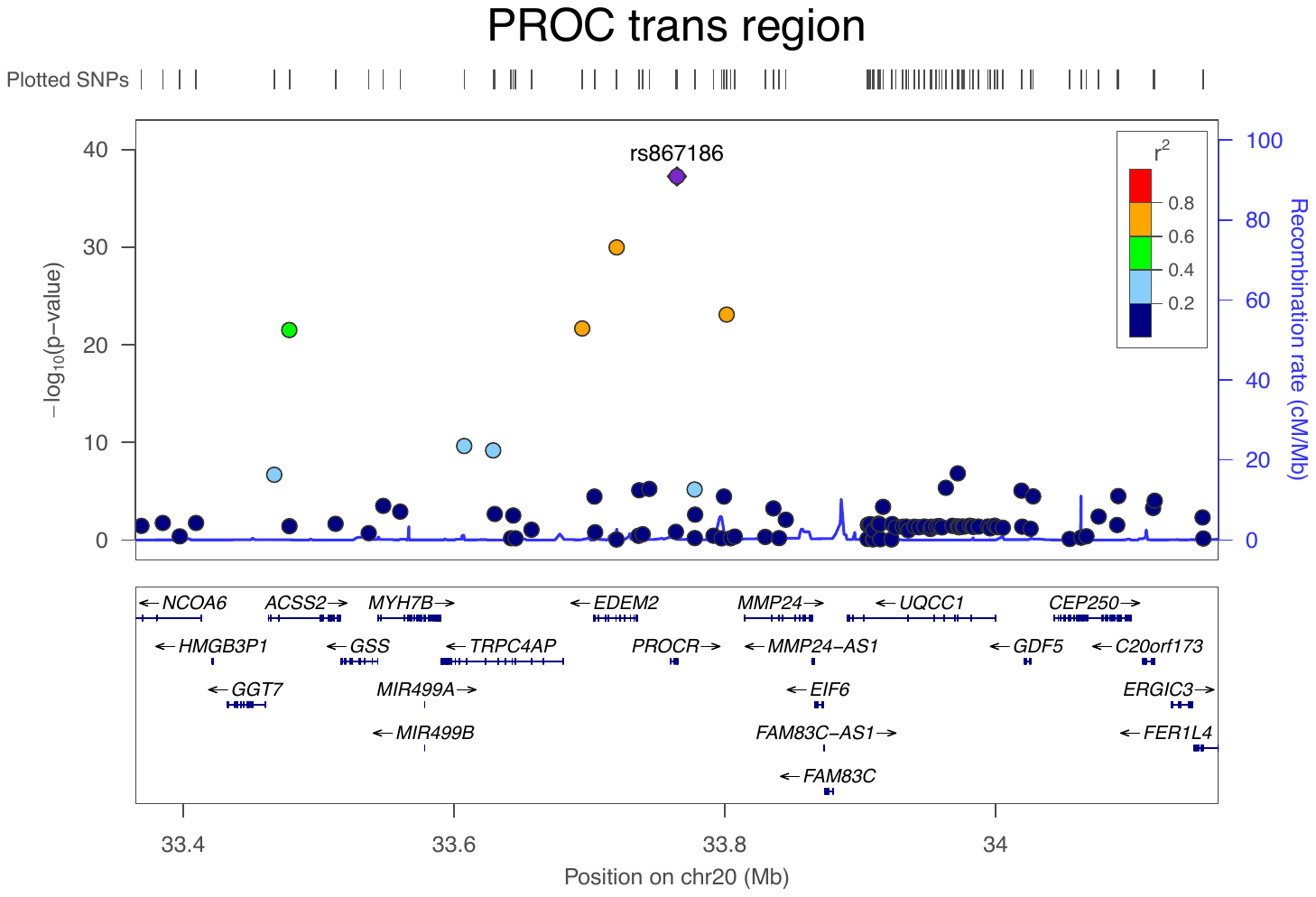

**Supplementary Figure 7.** Regional association plot of Protein C (PROC) protein levels in two regions. (A) there is little evidence that SNPs associated is with PROC protein levels in the cis region. (B) there is strong evidence that SNPs are associated with PROC in the trans PROCR region. The PROC data is from Suhre et al.

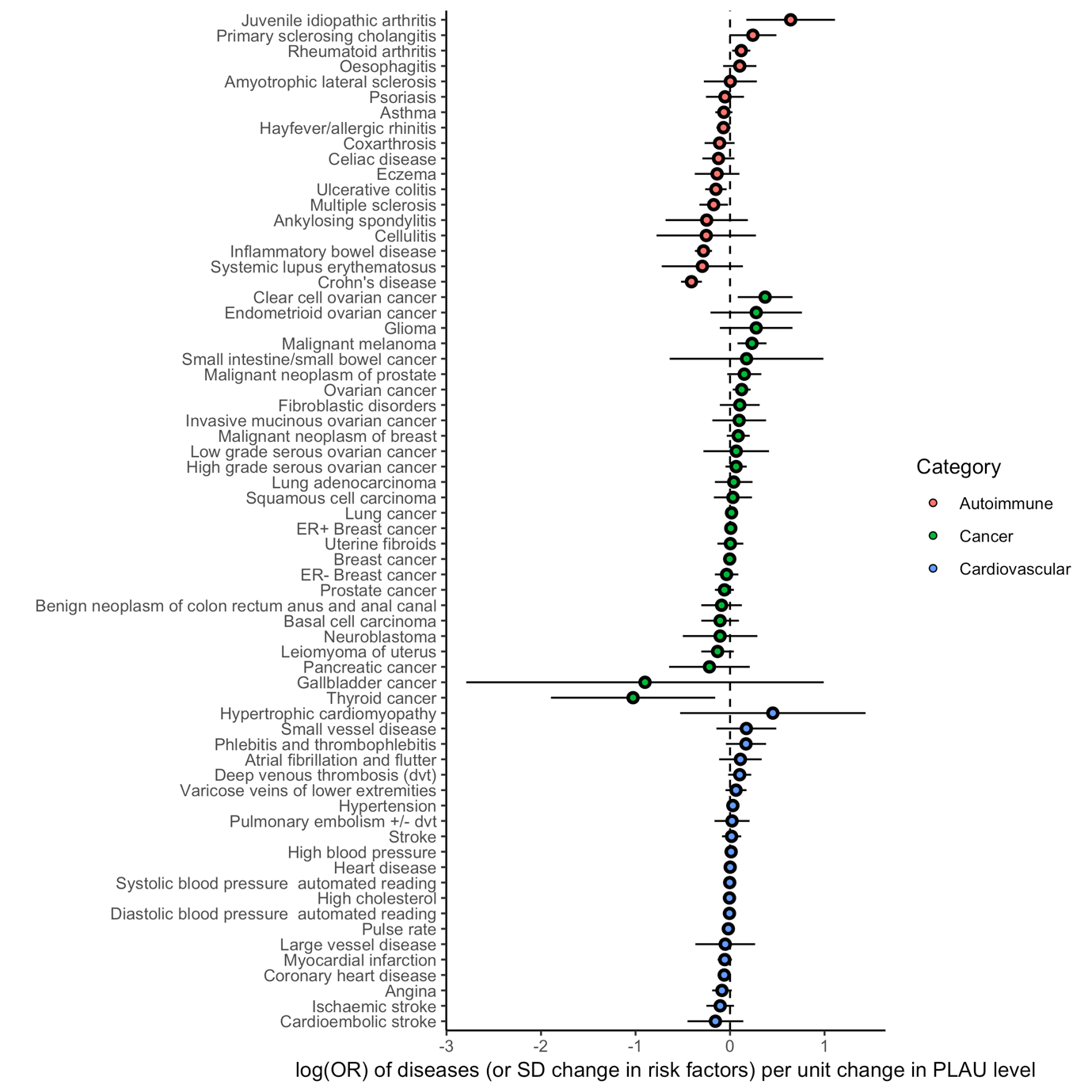

**Supplementary Figure 8.** Forest plot of MR estimates for plasma PLAU levels on immune-mediated phenotypes, cardiovascular phenotypes and cancers. The X-axis refers the log odds ratio of disease (or SD change in risk factor) per unit change in plasma PLAU level. The Y-axis refers to phenotypes tested in this Phenome-wide MR. The error bar refers to the 95% confidence interval of the causal estimates. Colours refer to different categories of phenotypes.

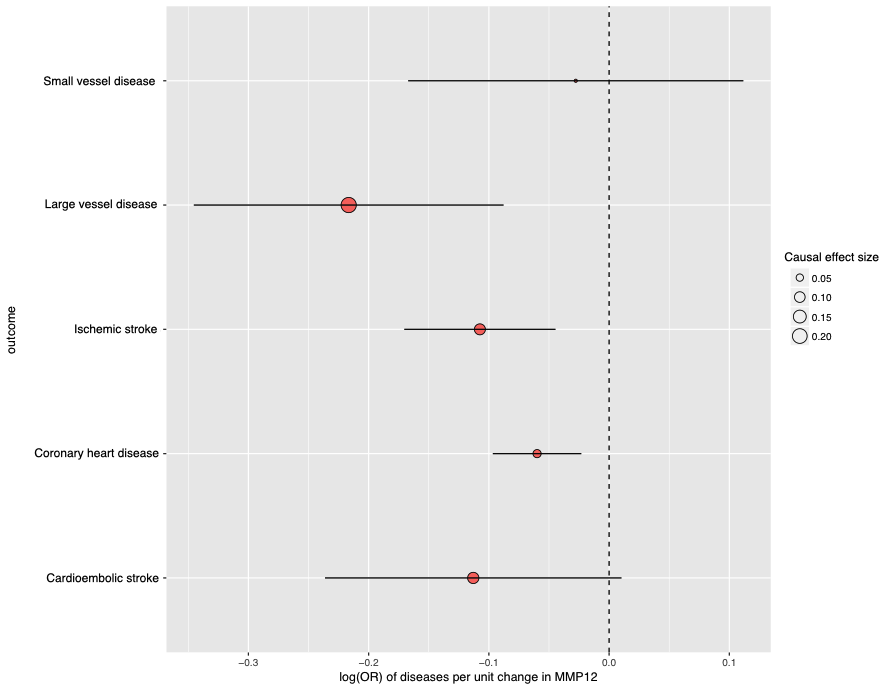

**Supplementary Figure 9.** Forest plot showing MMP12 MR associations on coronary heart disease and stroke. The X-axis refers the log odds ratio of disease per unit higher MMP12 level. The Y-axis refers to different subtypes of cardiovascular diseases. The error bar refers to the 95% confidence interval of the causal estimates.

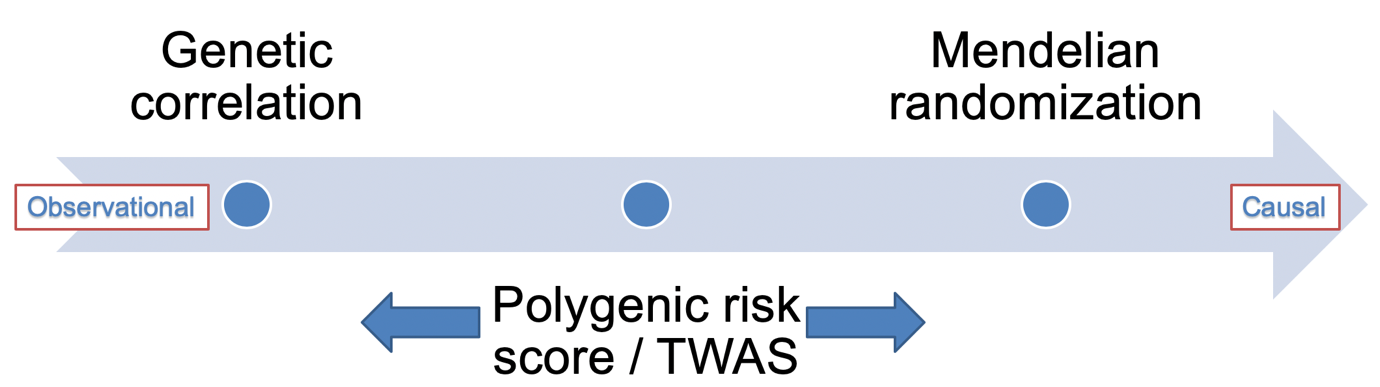

**Supplementary Figure 10.** Difference between Mendelian randomization, polygenic risk score / TWAS and genetic correlation. Note: Genetic correlation estimate the overall genetic overlap of SNPs across the whole genome, which the estimate is closer to observational correlation between two phenotypes with no direction. Mendelian randomization estimates the causal relationship between two phenotypes with a direction, which typically use SNPs robustly associated with the exposure (e.g. protein level) as instruments, where the instruments can be in both cis and trans region. The SNP effect on human phenotype will be estimated first and then meta-analysed using various models such as inverse variance weighted (IVW). The polygenic risk score association and TWAS are similar, which both need individual level genotype data. The method contains two steps, 1) using SNPs within the cis region to predict the expression of a gene; 2) correlate the predicted expression of the gene with the human phenotype.

#### Supplementary Note 1. Review of protein-trait associations in three disease areas

We found that our MR findings were clustered in three areas that have not been described well in the MR literature: blood pressure (AGT, ADM, ERAP2, FN1, SWAP70, CXCL16 and IGFBP3), lung function (ADAM19, APOF, GPC5, SERPINF1, MFAP2) and immune mediated disease (IL23R, IL6R, IL18R1, FCRL3, ICAM5 and PLAU).

For blood pressure, Adrenomedullin (ADM) is a hormone involved in vascular tone, and is a well-studied target for blood pressure ^1^ ^2^ *Endoplasmic Reticulum Aminopeptidase 2* (ERAP2) has been linked to pre-eclampsia ^3^, for which there are currently no effective drugs. Rare mutations in *Fibronectin 1* (FN1) have been associated with glomerulopathy with fibronectin deposits ^4^, which leads to hypertension ^5^. *C-X-C Motif Chemokine Ligand 16* (CXCL16) is an interferon-γ-regulated chemokine and scavenger receptor for oxidized low-density lipoprotein that is expressed in atherosclerotic lesions ^6^ and has been linked to Chronic Kidney Disease ^7^. These connections may suggest the association of CXCL16 on DBP is at least in part, through the atherosclerosis or renal function. Insulin-like growth factor binding protein 3 (IGFBP3) levels have been previously reported to associate with hypertension ^8^, and a SNP in IGFBP3 has been found to associate with increased long-term average pulse blood pressures ^9^.

For lung function, *A Disintegrin And Metalloproteinase Domain 19* (ADAM19) is a metalloproteinase, which may have a role in tissue remodelling ^10^. *Apolipoprotein F* (APOF) is a lipoprotein, which may associate with chronic obstructive pulmonary disease (COPD) ^11^. *Glypican Proteoglycan 5* (GPC5) is a cell surface heparin sulfate proteoglycan) that has been linked to lung cancer ^12^. *Serpin Family F Member 1* (SERPINF1) inhibits angiogenesis ^13^ and rare mutations are associated with osteogenesis imperfecta ^14^, but a link to lung function is unclear. *Microfibril Associated Protein 2* (MFAP2) is an antigen of elastin associated fibrils so may be important in tissue remodelling in the lung. However, the MFAP2 locus was associated with both height and lung function ^15^. It is possible that the association we identified could be mediated by height.

For the immune mediated traits, an existing IL23R antagonist, ustekinumab, demonstrated efficacy in reducing Crohn’s diseases in a recent phase III clinical trial ([www.clinicaltrials.gov](http://www.clinicaltrials.gov)) ^16^. Our MR analysis further linked IL23R inhibition with psoriasis. IL18R1 is a known eczema locus which replicated in both the Japanese and European population ^17^ ^18^. Our MR study suggested that IL18R1 could be an effector / causal gene for eczema. Polymorphisms within *Fc Receptor Like 3* (FCRL3) have been found to associate with rheumatoid arthritis in the Chinese population ^19^. A link between *Intercellular Adhesion Molecule 5* (ICAM5) and Crohn's disease is not yet clear in the literature.

#### Supplementary Note 2. Case study for our MR analysis replicated previous findings

Some of our MR results replicated those previously described by others. For example, Sun *et al* suggested that the TNFRSF11A associated variant rs884205 was also associated with Paget’s disease ^20^. Our Wald ratio analysis confirmed the positive association between this pQTL and Paget’s disease (OR=8.56, 95%CI=4.78 to 15.31, P= 4.72x10^-13^). The colocalization analysis further confirmed that the two traits share the same casual variant within the TNFRSF11A region (PP=99%).

In addition, we replicated the apparent effect of MMP12 pQTL on CHD (OR=0.94, 95%CI=0.91 to 0.98, P= 0.0014) and stroke reported in Sun *et al* ^20^. We extended this analysis to stroke subtypes and found that MMP12 pQTL were associated with ischemic stroke (OR=0.90, 95%CI=0.84 to 0.96, P=0.0008) and large vessel disease (OR=0.81, 95%CI=0.71 to 0.92, P=0.00098) but not strongly with cardioembolic stroke (OR=0.89, 95%CI=0.79 to 1.01, P=0.07) and not with small vessel disease (OR=0.97, 95%CI=0.85 to 1.12, P=0.70) (**Supplementary Table 26**, **Supplementary Figure 9**).

#### Supplementary Note 3. Detailed results of bi-directional MR and Steiger filtering for orienting causal direction in protein-disease associations

For the bi-directional MR, we modelled complex traits as our exposure (data from MR-Base) and plasma protein level as our outcome (full summary statistics of proteins were available for Sun *et al* and Folkersen *et al* ^20^*^,^*^21^). For clarity – this analysis does not necessarily implicate the disease status as being causal for the protein levels, but it indicates that genetic liability to the disease may influence protein levels. In total, the relationship between 104 diseases and 206 proteins were tested (841 tests, Bonferroni-adjusted P = 5.9x10^-5^). We found no strong evidence of reverse causality between protein level and disease for the majority of protein-trait associations (**Supplementary Data 1**). However, there were exceptions for traits with a strong genetic signal at the *APOE* locus.

Due to a lack of full summary statistics for some pQTL studies, we were not able to conduct bi-directional MR for all MR findings. Instead, we applied Steiger filtering as an alternative method to infer the causal direction for our MR. For MR findings using multiple instruments, the Steiger filtering tested the directionality of each instrument on exposure and outcome separately (rather than testing the overall directionality between exposure and outcome). Among all 397 protein-trait associations we tested, 360 had enough statistical power for the Steiger filtering analysis (to detect a nominally significant P value <0.05). For these 360 cases, 303 (84.2%) showed evidence that the direction of the association is from protein to human traits (**Supplementary Table 7, 8, 11, 12 and 13**). We found a substantially more reverse causal instances for trans (31.2%) than there are for cis (0.6%).

|  | Trans | Cis |
| --- | --- | --- |
| SF-FALSE | 53 | 1 |
| SF-TRUE | 117 | 178 |
| All | 170 | 179 |
| Percentage | 31.2% | 0.6% |

Note: SF-FALSE refers to protein-phenotype association with evidence of reverse causality using Steiger filtering; SF-TRUE means the association with no evidence of reverse causality.

##### Supplementary Note 4. Integration of protein-disease associations with gene expression evidence

To further strengthen the evidence of the protein-trait association, we integrated our pQTL MR results with gene expression evidence for a case study IL18R1 on Crohn’s disease. IL18R1 has two conditionally independent association signals in the cis region (top hit rs1420106 and second hit rs13014644). We investigated whether both cis pQTLs were also associated with gene expression of IL18R1 in various tissues and immune response stages. This was undertaken by querying data from 3 sources: the tissue specific eQTL data from GTEX ^22^ (sample size (n) from 80 to 491 for 48 tissues), whole blood eQTL data from eQTLGen ^23^ (n=31,684) and immune response QTL data from Kim-Hellmuth *et al.* ^24^ (n=134) (**Supplementary Table 18**). Evidence of eQTL association was defined as P < 1x10^-4^.

To evaluate whether the IL18R1-Crohn’s disease association was influenced by transcriptional level, we cross-referenced the two IL18R1 cis pQTLs (rs1420106 and rs13014644) with gene expression data from the GTEX ^22^ (sample size (n) from 80 to 491 for 48 tissues), the eQTLGen ^23^ (n=31,684) and Kim-Hellmuth *et al.* ^24^ (n=134) (**Supplementary Table 18**). The top hit, rs1420106, was associated with the expression level of *IL18R1* in 16 tissues (**Supplementary Table 18A**) and the SNP only influenced the immune response of IL18R1 under immune stimulation stage and would not be discovered in baseline monocytes (**Supplementary Table 18B**). In contrast, the second hit, rs13014644, showed much weaker influence in all immune response stages and only associated with expression level of IL18R1 in blood (**Supplementary Table 18A** and **B**).

#### Supplementary Note 5. The protocol of the instrument validation

Because instruments were identified from five independent studies performed using different analytical platforms (Box 1), we developed a protocol for instrument validation. **Figure 1** summarises the 2 key analyses used for instrument validation.

| Study | Platform | Sample size | Number of proteins (with pQTLs) | Number of instruments (pQTLs) |
| --- | --- | --- | --- | --- |
| Sun *et al* | SOMAScan | 3301 | 1478 | 1981 |
| Emilsson *et al* | SOMAScan | 3200 | 776 | 875 |
| Suhre *et al* | SOMAScan | 1000 | 284 | 539 |
| Folkersen *et al* | Olink | 3394 | 58 | 80 |
| Yao *et al* | xMAP | 6861 | 60 | 131 |

Box 1. The study level information of the 5 pQTLs studies.

##### 1 Harmonisation of Protein IDs and instruments

Since the 5 previous studies measured plasma protein levels using different probes from three different platforms (SOMAScan, Olink and Luminex xMAP), we mapped the platform ID for each protein analyte from each study to Uniprot IDs (and associated annotations) based on annotations provided by the platform vendors and manual review. We then grouped the analytes based on their Uniprot IDs. The end product included columns corresponding to the ‘platform ID’ of each the analyte from each study, the full protein name, gene symbol, Uniprot ID, Ensembl gene id and GRch38 gene location (**Supplementary Table 27**). Since a single probe from any platform can map to multiple proteins, each of the protein names, gene symbols, Uniprot IDs, Ensembl IDs and gene locations is given as a semicolon-delimited list of entries. In particular, we included all UniProt IDs for a given gene, which is essential for a robust mapping. Finally, using the platform IDs as a key, we collated the association information of the instruments from each study based on the Uniprot IDs (**Supplementary Table 27**).

##### 2 Instrument validation

##### 2.1. Combining and reassigning instruments to proteins

Since we regrouped the protein IDs based on their functions, we further reassigned the instruments to fit the new protein naming system rather than the protein names reported in the pQTL studies. We standardized the format of the instrument files across studies and reassigned them based on their Uniprot ID.

##### 2.2. Instrument specificity

Absence of horizontal pleiotropy is one of the core assumptions for MR, which assumes that the genetic variant should only be related to the outcome of interest through the instrumented exposure. We noted that some SNPs were associated with more than one protein, for example, APOE SNP rs7412 is associated with a set of proteins such as ADAM11, APBB2 and APOB. We considered these instruments associated with more than 5 proteins as potentially pleiotropic SNPs and non-specific for any particular protein level and set up a flag (discrete numeric parameter) based on the number of proteins these SNPs (and their proxies with LD r2>0.5) associated with (“N_protein” column in **Supplementary Table 1**).

##### 2.3. Instrument consistency across studies

##### 2.3.1. Cross-referencing association results between studies

Firstly, we identified all reported pQTLs across studies and harmonized the effects of SNP association so that the effect of a SNP on the protein and the effect of that SNP on the outcome corresponded to the same allele. We then looked up the results for the pQTLs in the other studies (e.g. for a pQTL reported by Sun et al, we looked up its SNP effect in Suhre et al). As the full GWAS summary statistics were not available for Yao et al, only 9 pair-wise lookups from 12 possible combinations could be conducted: validation could therefore be conducted between Sun pQTLs in Suhre and Folkersen; Suhre pQTLs in Sun and Folkersen; Folkersen pQTLs in Sun and Suhre, Yao pQTLs in Sun, Suhre and Folkersen. Among the 2113 SNPs, we found 1062 with effects in one or more other studies (**Supplementary Table 15**). Results of the pair-wise comparisons and the number of SNPs included in each comparison can be found in **Supplementary Table 2**.

##### 2.3.2. Pair-wise instrument validation

We noted some examples where SNPs were reported to be associated with a protein in one study but not reached the genome-wide p value threshold in other studies which had measured the same protein. In these instances, we investigated whether this reflected a no statistical evidence of association (in which case, this inconsistency may indicate potentially artefactual associations) or simply fluctuation of association strength, but with directionally consistent and nominally significant (p<0.05) associations in both studies (which would provide supporting evidence for an instrument).

Because of the low number of proteins measured in Folkersen *et al* and Yao *et al*, the number of cases where we could perform a validation across 4 or even 3 studies were limited. Instead, for the 1062 pQTLs with SNP lookup results in at least 2 studies, we performed 9 pair-wise comparisons to assess the consistency of the SNP effects of instruments in each pair of studies. Firstly, we tested the overall agreement of effect estimates for the pairwise comparisons. We estimated the pair-wise correlation (r), 95% confidence intervals and p values using the “cor.test” function in R (https://www.r-project.org/).

To provide an overall visualisation of the agreements we generated scatter plots for all pair-wise combinations (**Supplementary Figure 1**). For each scatter plot, we compared the genetic associations (betas, 95% CIs) from one study with the same SNPs looked up in the other study. We also generated Bland-Altman plots to compare the genetic effects between different studies (**Supplementary Figure 2**).

In summary, we show that the agreement of pQTL regression coefficients is high across all five studies (correlation ranged from r = 0.58 to 0.94, **Supplementary Table 2**), and in general, the effects of the SNP associations for all nine study-level pair-wise comparisons follows the identity line well with few outliers (**Supplementary Figure 1 and 2**). Furthermore, we defined three flags (binary parameter) based on 1) whether the direction of the effects agreed across studies (column “Agree_beta” in **Supplementary Table 15**); 2) whether P values of the SNP association were smaller than 0.05 across studies (column “Agree_P” in **Supplementary Table 15**); 3) whether there was statistical evidence of heterogeneity between the two effect sizes (pair-wise Z test, with Z score greater than 5 as threshold, which is equal to p value of 0.001) (column “Heterogeneity_across_studies” in **Supplementary Table 1**).

##### 2.3.3. Instrument validation using colocalization analysis across protein studies

For 82 instruments with SNP association information in both Sun *et al* and Folkersen *et al*, we further validated the instruments using a stringent Bayesian model implemented in “coloc” R package ^25^ to estimate the posterior probability (PP) of each genomic locus containing a single variant affecting the same protein in both Sun *et al* and Folkersen *et al* ^20^ ^26^. We set up a flag (binary parameter) to record SNPs colocalized in both studies (column “Coloc_across_studies” in **Supplementary Table 1**).

#### Supplementary Note 6. Description of ALSPAC study

Pregnant women resident in Avon, UK with expected dates of delivery 1st April 1991 to 31st December 1992 were invited to take part in the study. The initial number of pregnancies enrolled is 14,541 (for these at least one questionnaire has been returned or a “Children in Focus” clinic had been attended by 19/07/99). Of these initial pregnancies, there was a total of 14,676 foetuses, resulting in 14,062 live births and 13,988 children who were alive at 1 year of age.

When the oldest children were approximately 7 years of age, an attempt was made to bolster the initial sample with eligible cases who had failed to join the study originally. As a result, when considering variables collected from the age of seven onwards (and potentially abstracted from obstetric notes) there are data available for more than the 14,541 pregnancies mentioned above.

The number of **new pregnancies** not in the initial sample (known as Phase I enrolment) that are currently represented on the built files and reflecting enrolment status at the age of 24 is 904 (452, 254 and 198 recruited during Phases II, III and IV respectively), resulting in an additional 811 children being enrolled. The phases of enrolment are described in more detail in the cohort profile paper (see footnote 4 below). Please note that phase 4 enrolment (age 18-24) is not currently included in the cohort profile.

The total sample size for analyses using any data collected after the age of seven is therefore 15,247 pregnancies, resulting in 15,458 foetuses. Of this **total sample** of 15,656 foetuses, 14,973 were **live births** and 14,899 were **alive at 1 year of age**.

A 10% sample of the ALSPAC cohort, known as the **Children in Focus (CiF) group**, attended clinics at the University of Bristol at various time intervals between 4 to 61 months of age. The CiF group were chosen at random from the last 6 months of ALSPAC births (1432 families attended at least one clinic). Excluded were those mothers who had moved out of the area or were lost to follow-up, and those partaking in another study of infant development in Avon.

Ethical approval for the study was obtained from the ALSPAC Ethics and Law Committee and the Local Research Ethics Committees.

Please note that the study website contains details of all the data that is available through a fully searchable data dictionary and variable search tool: http://www.bristol.ac.uk/alspac/researchers/our-data/
